# Injury-transduced SerpinOLs modulate neuroinflammation and glial activation in the diseased and non-diseased CNS

**DOI:** 10.64898/2026.01.29.702669

**Authors:** Yan Wang, Meina Zhu, Joohyun Park, Hyun Kyu Kim, Xiong Ge, Xi Chen, Bo Chen, Shinghua Ding, Wensheng Lin, Fuzheng Guo

## Abstract

Oligodendroglial dysfunction is common in CNS diseases and injuries, but the molecular and functional responses of oligodendroglia to CNS pathologies remain poorly defined. Here, we report that oligodendrocytes (OLs) respond to demyelinating diseases by dysregulating serine protease inhibitor clade A member 3N (SERPINA3N). Homeostatic OLs transition to Serpina3n-expressing OLs (SerpinOLs) under other diseased conditions including stroke, endotoxicity, neurodegeneration, neurotrauma, and non-diseased healthy aging conditions. Mechanistically, general neuroinflammation or inflammatory mediators is insufficient for SerpinOL transition. Instead, oligodendrocyte damage/injury, even in the absence of neuroinflammation or glial activation, is sufficient for the transition. Phenotypically, SerpinOLs are characterized by molecular signatures of inflammatory and immune regulation and STAT3 signaling activation. Functionally, SerpinOLs exacerbate neuroinflammation and promote glial activation toward pro-inflammatory and neurodegenerative states. Together, our findings suggest SerpinOLs are a common population of injury-transduced OLs that amplify neuroinflammation and glial activation in the diseased and non-diseased CNS through SERPINA3N secretion. Our findings provide new insights into myelination-independent role of OLs in regulating CNS pathophysiology.

## Introduction

Oligodendrocytes (OLs), generated from oligodendroglial progenitor cells (OPCs), are myelin-forming cells of the central nervous system (CNS). Oligodendroglial lineage cells (OLs and OPCs) support neuron/axonal survival and shape neural circuitry through developmental myelination in the developing brain ^1^ and adaptive myelination in the adult brain ^2,3^, both of which are essential for brain function and behavior ^4–6^. OLs and their myelin derivatives are the primary victims of demyelinating disorders such as multiple sclerosis, an inflammatory CNS demyelinating disease. In addition, oligodendrocyte dysfunction has been well-recognized in other CNS pathologies ^7^ such as stroke ^8,9^, endotoxicity ^10,11^, neurodegeneration^12,13^, neurotrauma ^14^, and even normal (or healthy) aging, a naturally occurring process in the absence of diseases or trauma ^15,16^. Despite the commonly existing dysfunction, how OLs respond to CNS pathologies molecularly and functionally remains incompletely understood.

Single-cell transcriptomics and meta-analysis have unveiled diverse subpopulations of disease-associated or specific oligodendroglia (DAO)^17–21^ in the diseased CNS. They are defined by different transcriptomic signatures, for example, DAO defined by signature genes associated with immunogenesis, differentiation and survival, and interferon-response^18,22^. There is no doubt that more DAO subpopulations will be identified with the technical advancement of the next-generation sequencing and bioinformatics. However, there are at least two fundamental knowledge gaps for the ever-growing DAO subpopulations. First, the cellular and molecular mechanisms that transition homeostatic OLs into distinct activation states are still enigmatic. Second, the biological functions of DAO in regulating disease pathophysiology remain unknown or largely speculative *in vivo*^17,23–25^.

In the present study, we found that homeostatic OLs respond to inflammatory demyelination by secreting serine protease inhibitor clade A member 3N (SERPINA3N), a secretory protein dysregulated in the brain and body fluids in many neurological conditions ^26^. SERPINA3N-expressing OLs (termed as SerpinOLs in the study) are present in various types of CNS neurological conditions. Using a series of transgenic mice and injury models, our findings suggest that direct intrinsic injury to OLs, regardless of the presence of neuroinflammation (a common feature of CNS pathologies), triggers the conversion of homeostatic OLs into SerpinOLs. Using SERPINA3N deficiency mice, we demonstrate that SerpinOLs perpetuate CNS inflammatory response and promote glial activation toward pro-inflammatory and neurodegenerative states in the diseased and aged CNS. Thus, our study defines SerpinOLs as an injury-transduced population of OLs that serve as crucial perpetuators for neuroinflammation and glial activation, common pathological features in various neurological diseases and injuries. Our findings indicate OLs actively participate in CNS pathology regulation through myelination-independent pathways.

## Results

### Screening for demyelination-responsive genes identifies *Serpina3n* dysregulation selectively in oligodendrocytes

We set off to identify genes responsive to inflammatory demyelination by conducting an unbiased RNA-seq analysis of MOG/EAE mice (**Suppl Fig. 1a**), an animal model of multiple sclerosis^27^. Most top differentially expressed genes (DEGs), including *Gpnmb*, *Trem2*, *Lyz2*, *Ctsd*, and *Cd68*, were enriched in microglia/macrophages (**Suppl Fig. 1b**). In contrast, *Serpina3n,* the third most highly expressed DEG in the top 10 list (**Suppl Table 1**), was selectively induced in oligodendrocytes (OLs) (**Fig. 1a**), as shown by glial cell type-specific RNA-seq analysis (**Suppl Fig. 1c-f**). Among OL-specific DEGs, *Serpina3n* showed the strongest induction in response to MOG/EAE (**Suppl Table 2**), a result confirmed by real-time quantitative PCR (RT-qPCR) (**Fig. 1b**). These findings identify *Serpina3n* as a responsive gene to inflammatory demyelination specifically in OLs.

**Figure 1.**
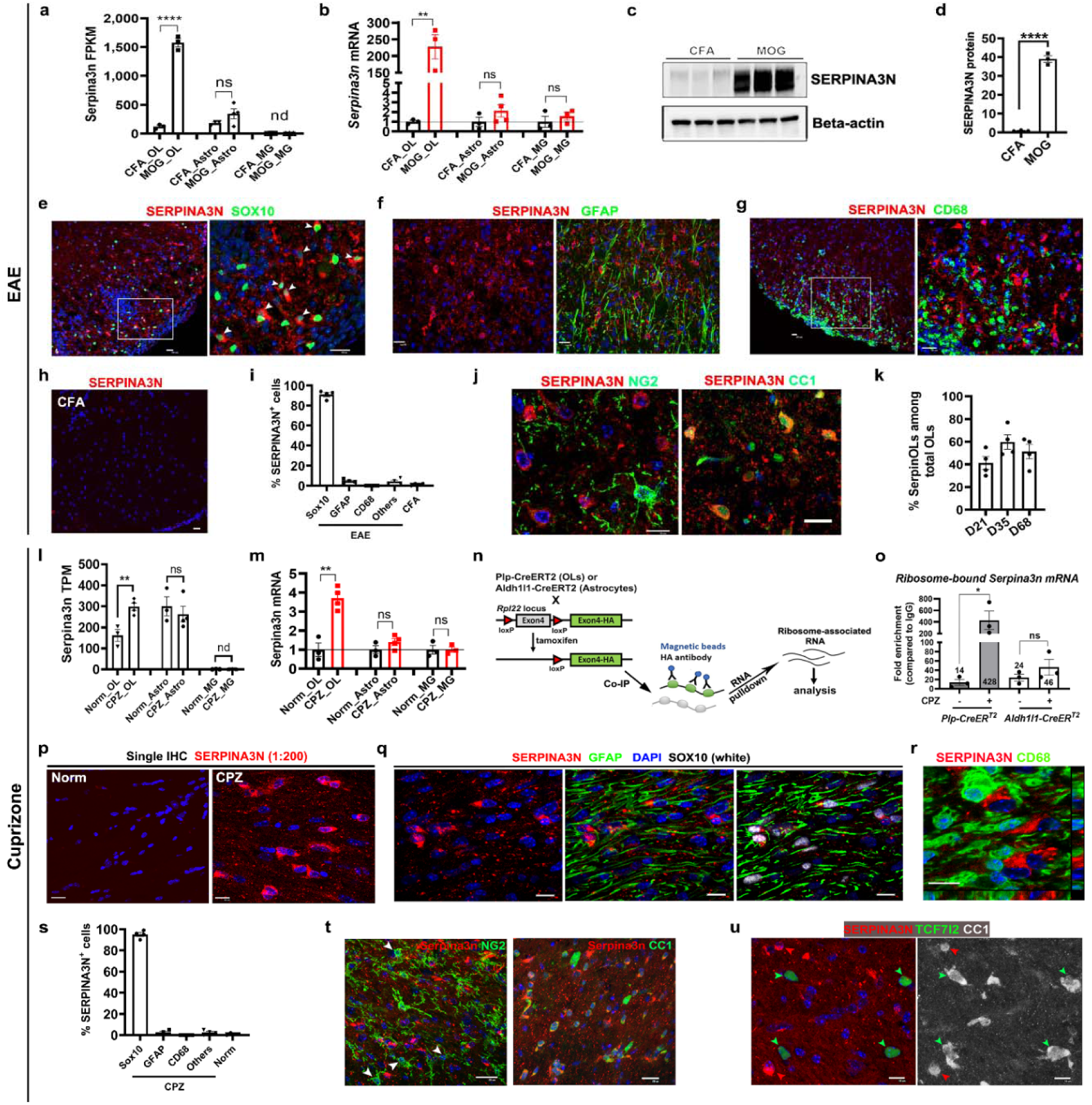
Oligodendrocytes transition into SERPINA3N-expressing OLs (SerpinOLs) in response to CNS demyelination. **a-b** RNA-seq (**a**) and RT-qPCR (**b**) in MACS-purified oligodendroglia (OL), astroglia (Astro), and microglia (MG) from D30 MOG/EAE and CFA Ctrl spinal cords. nd, not detectable. (**a**), OL, *\*\*\*\* P* < 0.0001; Astro, *P* = 0.1684; MG, *P* = 0.2453. (**b**), OL, *\*\* P* = 0.0035; Astro, *P* = 0.2404; MG, *P* = 0.4565. *n* = 3 mice except for MOG_Astro (n=4). **c-d** Western blot (**c**) and quantification (**d**) in D21 MOG/EAE and CFA spinal cords. *\*\*\*\* P* < 0.0001. *n* = 3 mice. **e-i** Immunofluorescence (IHC) for SERPINA3N with SOX10 (**e**), GFAP (**f**) or CD68 (**g**) in D21 MOG/EAE and CFA Ctrl (**h**) spinal cord ventral white matter and % SERPINA3N⁺ cells by lineage markers (**i**). Arrowheads, intracellular SOX10⁺SERPINA3N⁺ cells. Boxed areas shown in higher magnification. *n* = 4 mice. **j-k** IHC of SERPINA3N and NG2 (OPC marker) or CC1 (mature OL marker) in D21 MOG/EAE spinal cords (**j**) and quantification of SERPINA3N^+^CC1^+^ SerpinOLs among CC1^+^ OLs (**k**). *n* = 4 mice each time-point. **l-m** RNA-seq (**l**) and RT-qPCR (**m**) in MACS-purified OL, Astro and MG from brains of 4-week cuprizone (CPZ) or normal (Norm) diet mice. (**l**) OL, *\*\* P* = 0.0075; Astro, *P* = 0.5503; MG, *P* = 0.6745. (**m**), OL, *\*\* P* = 0.0019; Astro, *P* = 0.2039; MG, *P* = 0.9847. *n* = 3 mice Norm_OL, Norm_Astro, Norm_MG, CPZ_MG; *n* = 4 mice CPZ_OL, CPZ_Astro. **n** RiboTag strategy to analyze ribosome-bound transcripts in OL and Astro using *Plp1*-CreER^T^^2^ and *Aldh1l1*-CreER^T2^ Cre lines. Mice received tamoxifen (i.p., daily for 5 days), followed by 14 days clearance time prior to 4 weeks CPZ or normal diet. RNA was immunoprecipitated using anti-HA antibody or IgG Ctrl. **o** RT-qPCR of ribosome-bound *Serpina3n* transcripts. * *P* = 0.0427, ns, *P* = 0.9971. *n* = 3 mice each group. **p** IHC of SERPINA3N in corpus callosum (CC) of Norm (left) and 4-week CPZ diet (right). **q** Triple IHC of SERPINA3N, SOX10, and GFAP. **r** orthogonal view of confocal images showing absence of SERPINA3N in CD68^+^ cells. **s** % SERPINA3N^+^ cells co-expressing the indicated markers. *n* = 4 mice each group. **t** Double IHC of SERPINA3N and NG2 or CC1. Arrowheads, NG2^+^ OPCs. **u** Triple IHC for SERPINA3N, CC1, and newly regenerated OL marker TCF7l2. Green arrowheads, CC1^+^TCF7l2^+^SERPINA3N^-^ cells; red arrowheads, CC1^+^TCF7l2^-^SERPINA3N^+^ cells. **p-u**, IHC was imaged in the corpus callosum after 4-week CPZ (or Norm) diet. Scale bars: **e-h, j, p, q, r, u** 10 µm, **t** 20 µm. Data were presented as mean ± s.e.m. (a, b, d, i, k, l, m, o, s)

### SERPINA3N is dysregulated and secreted by oligodendrocytes during autoimmune demyelination

To determine whether *Serpina3n* mRNA induction results in protein upregulation, we performed Western blot (**Fig. 1c**), which revealed a >30-fold increase in SERPINA3N protein in the spinal cord of MOG/EAE mice (**Fig. 1d**). Fluorescent immunohistochemistry (IHC) was used to characterize the cellular specificity of SERPINA3N. Not only intracellular but also extracellular SERPINA3N was detected in MOG/EAE spinal cord (**Fig. 1e-g**), consistent with its secretory properties ^26^. SERPINA3N^+^ cells were identified as SOX10^+^ oligodendroglial lineage cells (**Fig. 1e, i**) and few were GFAP^+^ astrocytes (**Fig. 1f, i**) or CD68^+^ activated microglia/macrophages (**Fig. 1g, i**). In CFA-treated control mice, SERPINA3N expression was minimal (**Fig. 1h, i**). All SERPINA3N^+^ oligodendroglial lineage cells were identified as CC1^+^ mature OLs but not NG2^+^ OPCs (**Fig. 1j**). These findings suggest that OLs upregulate and secrete SERPINA3N in response to inflammatory demyelination.

SERPINA3N-expressing OLs, which we referred to as SerpinOLs, persisted into the chronic phase of MOG/EAE (**Suppl Fig. 2a-c**) when peripheral inflammatory infiltrates subside ^28^, indicating that peripheral immune cell infiltration is dispensable for SERPINA3N expression. SerpinOLs were found in both lesional (**Suppl Fig. 2b**) and non-lesional regions (**Suppl Fig. 2c**). SerpinOLs were negative for TCF7l2 (**Suppl Fig. 2c**), a nuclear marker labeling newly regenerated OLs^29^, suggesting that they are not newly formed OLs. Many SerpinOLs were located in proximity to CD68^+^ cells (**Suppl Fig. 2d**). At the population level, SerpinOLs accounted for ∼40-60% of all mature OLs during the disease course of MOG/EAE (**Fig. 1k**). Together, our data suggest that OLs, but not other cell types, respond to inflammatory demyelination via SERPINA3N induction.

### Chemical-induced demyelination transitions homeostatic OLs into SerpinOLs

We next tested whether OLs are the major cell type secreting SERPINA3N in the cuprizone (CPZ) demyelination model, which was characterized by the local expansion and activation of resident microglia with limited number of monocyte-derived macrophages (Ccr2-RFP^+^) (**Suppl Fig. 3a**). Glial type-specific RNA-seq (**Suppl Fig. 3b-c, Suppl Table 3**) showed a significant increase of *Serpina3n* transcripts only in OLs (**Fig. 1l**), a finding validated by RT-qPCR assay (**Fig. 1m**) although astrocytes exhibited similar levels of *Serpina3n* transcripts to OLs (**Fig. 1l**).

We used the RiboTag technique ^30^ to determine if *Serpina3n* transcripts are translated into SERPINA3N proteins in astrocytes and/or OLs (**Fig. 1n**). Our results revealed >30-fold increase in ribosome-bound *Serpina3n* mRNA in OLs, but not astrocytes (**Fig. 1o**) during CPZ demyelination. This finding suggests that the active translation of SERPINA3N protein occurs predominantly in OLs during CPZ demyelination.

SERPINA3N showed a splenium-to-genu gradient in the corpus callosum of CPZ-treated mice (**Suppl Fig. 3d**), mirroring the caudal-to-rostral gradient of oligodendrocyte/myelin damage in this model ^31^. Both intracellular and extracellular SERPINA3N signal was present in CPZ-treated mice, whereas SERPINA3N was barely detectable in normal diet controls (**Fig. 1p**). Greater than 90% of intracellular SERPINA3N^+^ cells were positive for SOX10 and few, if any, GFAP (**Fig. 1q, s**) or CD68 (**Fig. 1r, s**), identifying them as oligodendroglial lineage cells. All SERPINA3N^+^ oligodendroglial lineage cells were further identified as CC1^+^ mature OLs. but not NG2^+^ OPCs (**Fig. 1t**) or TCF7l2^+^ newly generated OLs ^29^ (**Fig. 1u**). Paradoxically, extensive co-labeling of SERPINA3N with GFAP^+^ reactive astrocytes was noticed only on fluorescent double IHC of SERPINA3N/GFAP (**Suppl Fig. 3e**), an observation also reported in previous studies ^32,33^. The observed co-labeling likely resulted from optical artifacts because the process-like SERPINA3N immunoreactive signal (**Suppl Fig. 3e**) was absent from the single IHC of SERPINA3N (**Fig. 1p**), triple IHC of SERPINA3N/GFAP/SOX10 (**Fig. 1q, Suppl Fig. 3f**), or sequential staining of SERPINA3N and GFAP (and the reverse order) (**Suppl Fig. 3g-h**). Collectively, our results show that OLs upregulate and secrete SERPINA3N in response to CNS demyelination regardless of peripheral immune infiltration.

### Genetic evidence demonstrating oligodendrocyte-specific SERPINA3N upregulation and secretion during demyelinating injury

Due to its secretory nature, SERPINA3N in OLs may originate from other cell types and deposit into OLs. No direct evidence exists proving that SERPINA3N protein originates from OLs. To study the cellular origin of SERPINA3N, we generated *Serpina3n-tdTom* reporter mice (**Fig. 2a, Suppl Fig. 4a-d**). The induction of SERPINA3N-driven tdTom expression was validated using MOG/EAE and CPZ models (**Suppl Fig. 4e-f**). In MOG/EAE spinal cord, tdTom was co-localized with intracellular SERPINA3N (**Fig. 2b**, arrowheads), confirming the efficacy of tdTom in reporting SERPINA3N expression. Over 90% of tdTom^+^ cells were SOX10^+^ oligodendroglial cells with a neglectable fraction being GFAP^+^ astroglial cells or CD68^+^ microglia/macrophages (**Fig. 2c-e**). Similarly, in the corpus callosum of CPZ-demyelinating mice, >90% of tdTom^+^ cells were identified as SOX10^+^ oligodendroglial and few GFAP^+^ astroglial cells or CD68^+^ microglia (**Fig. 2f-h**), suggesting that OLs but not astrocytes are the cellular origin of CNS SERPINA3N protein. To further corroborate our conclusion, we generate oligodendroglia-specific Serpina3n conditional knockout (cKO) mice (*Olig2-Cre:Serpina3n*^fl/fl^) ^34^. In these mice, SERPINA3N immunoreactivity, both intracellular and extracellular, was nearly abolished in the spinal cord of MOG/EAE mice (**Fig. 2i**) and in the corpus callosum of CPZ mice (**Fig. 2j**). Thus, these data provide the first line of evidence conclusively demonstrating that OLs are the primary source of SERPINA3N expression and secretion in response to CNS demyelination.

**Figure 2.**
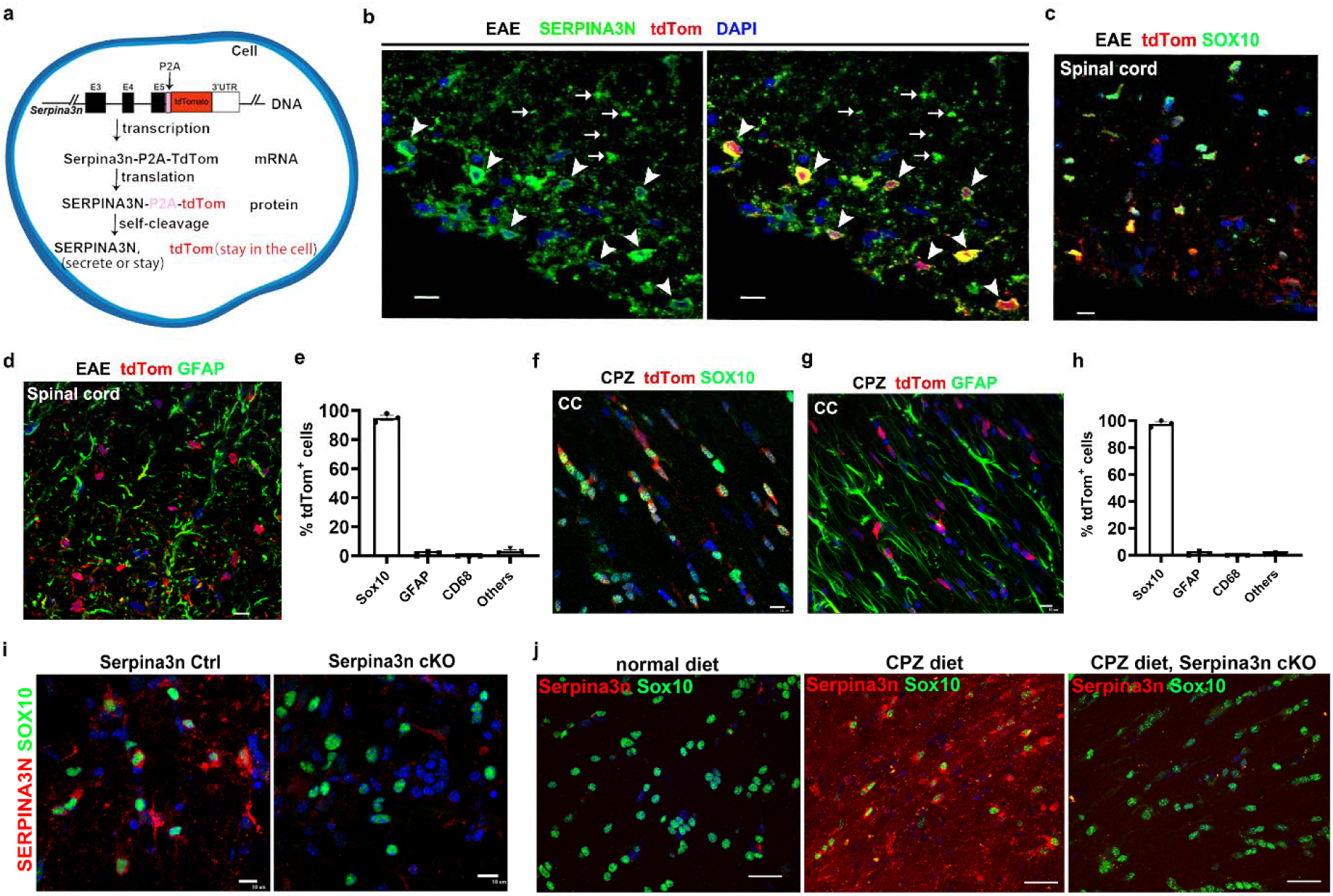
Genetic evidence for SERPINA3N expression in oligodendrocytes. **a** Schematic of Serpina3n-tdTom reporter design. The P2A self-cleaving site separates dTom from SERPINA3N-tdTom, allowing tdTom to localize in cell bodies while SERPINA3N remains intracellular or is secreted. **b** Double fluorescent IHC for SERPINA3N and tdTom in D30 MOG/EAE ventral white matter of spinal cord. Arrowheads, tdTom^+^ SERPINA3N^+^ cell bodies; arrows, extracellular SERPINA3N. **c-d** IHC of tdTom and SOX10 (**c**) or GFAP (**d**) in D30 MOG/EAE spinal cord. **e** Quantification of tdTom^+^ cells co-expressing indicated marker in D30 MOG/EAE spinal cord. *n* = 3 mice each group. **f-g** IHC of tdTom and SOX10 (**f**) or GFAP (**g**) in 3-week CPZ corpus callosum. **h** Quantification of tdTom^+^ cells co-expressing indicated markers in 3-week CPZ corpus callosum. *n* = 3 mice each group. **i** IHC for SERPINA3N and SOX10 in spinal cord of *Serpina3n* control (Ctrl) and conditional knockout (cKO, *Olig2-Cre:Serpina3n*^fl/fl^) mice at D30 post-MOG/EAE. **j** IHC of SERPINA3N and SOX10 in corpus callosum of *Serpina3n* Ctrl and cKO mice after 4-week CPZ (or Norm) diet. Scale bars: **b, c, d, f, g, i** 10 µm, **j** 20 µm. Data were presented as mean ± s.e.m. (e, h).

### Diverse CNS pathologies induce the transition of homeostatic OLs into SerpinOLs

Previous studies have identified *Serpina3n* as a marker of reactive astrocytes in diseased conditions such as ischemic stroke ^35^, lipopolysaccharide (LPS)-induced neuroinflammation ^35,36^, Alzheimer’s disease (AD)^37^, and non-diseased normal aging ^37,38^. However, our findings in demyelination models prompted us to re-evaluate this widely cited concept. We hypothesize that OLs are the primary source of SERPINA3N in these CNS pathologies.

In the ischemic stroke model induced by photothrombosis ^39^, neuronal damage/loss occurred acutely by 2 days post-injury (dpi) while glial scar formation became prominent in the chronic phase by 7 dpi (**Suppl Fig. 5 a1-a3**). We found that SERPINA3N^+^ cells peaked in the chronic phase by 7 dpi, predominantly in penumbra regions (**Suppl Fig. 5 b1-b2**). More than 70% of SERPINA3N^+^ cells were SOX10^+^ oligodendroglia in the penumbra and lesional regions at 7 dpi (**Fig. 3 a1, a3**) and 14 dpi (**Fig. 3 b1, b3**). In contrast, fewer than 10% were GFAP^+^ reactive astrocytes (**Fig. 3 a2-a3, Fig. 3 b2-b3, Suppl Fig. 5 c1-d2**) and none were CD68^+^ myeloid cells (**Suppl Fig. 5 e**) throughout the disease course. These results indicate that OLs are the main source of SERPINA3N in the ischemic brain, particularly during chronic stages.

**Figure 3.**
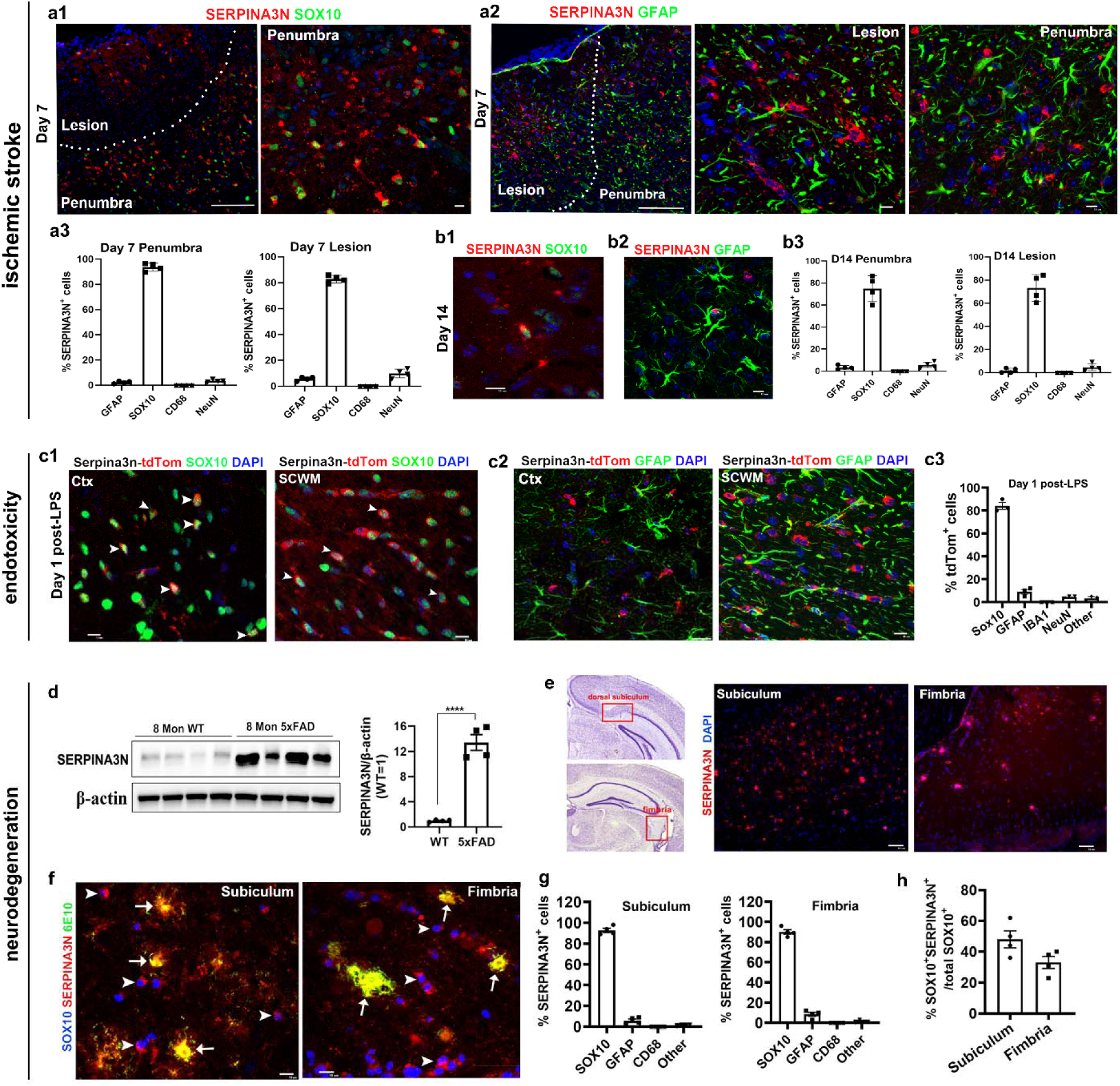
CNS insults transition homeostatic oligodendrocytes into SerpinOLs across various brain pathologies. **a1-a3**, fluorescent IHC of SERPINA3N with SOX10 or GFAP and quantification (n=4) at day 7 post-injury (dpi) of photothrombosis-induced stoke in mice. Scale bar: 100 µm (low magnification), 10 µm (high magnification). **b1-b3**, fluorescent IHC and quantification at 14 dpi. Scale bar: 10 µm. **c1-c3**, fluorescent IHC and quantification of tdTom (Serpina3n-tdTom reporter) and SOX10 or GFAP at D1 post-LPS injection. Ctx, cerebral cortex, SCWM, subcortical white matter. Scale bars: 10 µm. **d**, Western blot showing elevated SERPINA3N expression in brains of 5xFAD mice vs. age-matched control mice. *t*_(6)_ = 9.844 *P* < 0.0001. **e**, fluorescent IHC of SERPINA3N in the subiculum and fimbria of 7-8 Mon 5xFAD mice. Scale bars: 10µm. **f**, triple fluorescent IHC of SERPINA3N, SOX10, and β-amyloid (6E10) in subiculum and fimbria of 5xFAD mice. Arrows, co-localization of extracellular SERPINA3N with β-amyloid plaques; arrowheads, co-localization of intracellular SERPINA3N with SOX10. Scale bars: 10µm. **g**, percentage of SERPINA3N^+^ cells positive for indicated markers in 5xFAD mice. **h**, proportion of SOX10^+^ cells expressing SERPINA3N in 5xFAD mice. Data were presented as mean ± s.d. (a3, b3, c3, d) and mean ± s.e.m. (g, h).

In the LPS-induced neuroinflammation model, over 70% of SERPINA3N^+^ cells at 1 and 3 days post-LPS injection were SOX10^+^ oligodendroglia and <20% were GFAP^+^ or Aldh1l1-eGFP^+^ astrocytes and very few, if any, IBA1^+^ microglia or NeuN^+^ neurons (**Suppl Fig. 6 a1-a5**). To corroborate these findings, we utilized *Serpina3n-tdTom* transgenic mice. Approximately 70-80% of tdTom^+^ cells were SOX10^+^ oligodendroglial cells on day 1 (**Fig. 3 c1**, **c3**) and day 3 (**Suppl Fig. 6 b1, b3**) post-LPS injection. In contrast, < 20% of tdTom^+^ cells were GFAP^+^ astroglial cells in both time points (**Fig. 3 c2, c3, Suppl Fig. 6 b2, b3**). Thus, oligodendroglia, but astroglia, are the predominant cell types expressing SERPINA3N in response to LPS-elicited neuroinflammation.

In the 5xFAD mouse model, which recapitulates amyloid-beta (Aβ) pathology in AD patients, SERPINA3N expression was significantly upregulated in the brain of 7-8 months old mice (**Fig. 3d**) where it was found primarily in the hippocampal subiculum and fimbria (**Fig. 3e**). Large deposits of extracellular SERPINA3N were co-labeled with Aβ plaques (**Fig. 3f**, **Suppl Fig. 7a**) and surrounded by CD68^+^ activated microglia (**Suppl Fig. 7b-d**). Intriguingly, more than 90% of intracellular SERPINA3N^+^ cells were SOX10^+^ oligodendroglia (**Fig. 3g, Suppl Fig. 7e**). Quantification revealed that 30-50% oligodendroglia were transitioned into SerpinOLs in the subiculum and fimbria of 7-8-month-old 5xFAD mice (**Fig. 3h**). Paradoxically, double fluorescent IHC showed a nearly 100% overlap between SERPINA3N immunoreactive signal and GFAP^+^ reactive astrocytes in the hippocampus (**Suppl Fig. 8a**) and subiculum (**Suppl Fig. 8c**). This phenomenon, which was also reported in previous studies ^37^, is likely resulted from optical artifacts, as the observed astrocyte-like SERPINA3N signal was not seen in single IHC of SERPINA3N (**Suppl Fig. 8b, d**), double IHC of SERPINA3N/SOX10 (**Suppl Fig. 7e**), or triple IHC of SERPINA3N/GFAP/SOX10 (**Suppl Fig. 8e**), indicating optical artifacts. Thus, we conclude that OLs are the major cell types that respond to AD-like amyloidosis by dysregulating SERPINA3N expression.

In the murine spinal cord injury model elicited by lateral hemi-sectioning (**Suppl Fig. 9a**), SERPINA3N^+^ cells were localized in the white matter below the injured spinal segments (**Suppl Fig. 9b-c**). Nearly all were SOX10^+^ oligodendroglia, with few being GFAP^+^ astroglia or CD68^+^ microglia (**Suppl Fig. 9d-f**). These data suggest that OLs but not astrocytes or microglia are the major cell population producing SERPINA3N during neurotrauma.

Taken all together, our findings demonstrate that homeostatic OLs transition into SerpinOLs in response to various neurological diseases and injuries. These results also challenge the widely cited concept that Serpina3n is a reliable marker of reactive astrocytes in the diseased CNS ^35^.

### Normal aging transitions homeostatic OLs into SerpinOLs

We next examine whether normal healthy aging, a naturally occurring process absent of diseases or neurotraumas ^16,40^, is sufficient to convert homeostatic OLs into SerpinOLs. While homeostatic OLs do not express SERPINA3N in the young adult brain (**Fig. 4a**), we observed robust induction of intracellular and extracellular SERPINA3N in the aged brain (20 months old), predominantly in the subcortical white matter tracts, such as corpus callosum (**Fig. 4b**) and fimbria (**Fig. 4d**), consistent with recent findings that white matter is the most vulnerable “hotspot” of normal aging^41,42^. Approximately 90% of intracellular SERPINA3N^+^ cells were identified as SOX10^+^ oligodendroglia (**Fig. 4b, d**, arrowheads, **Fig. 4i**). Further analysis confirmed that they were CC1^+^ mature OLs (**Fig. 4f**, arrowheads).

**Figure 4.**
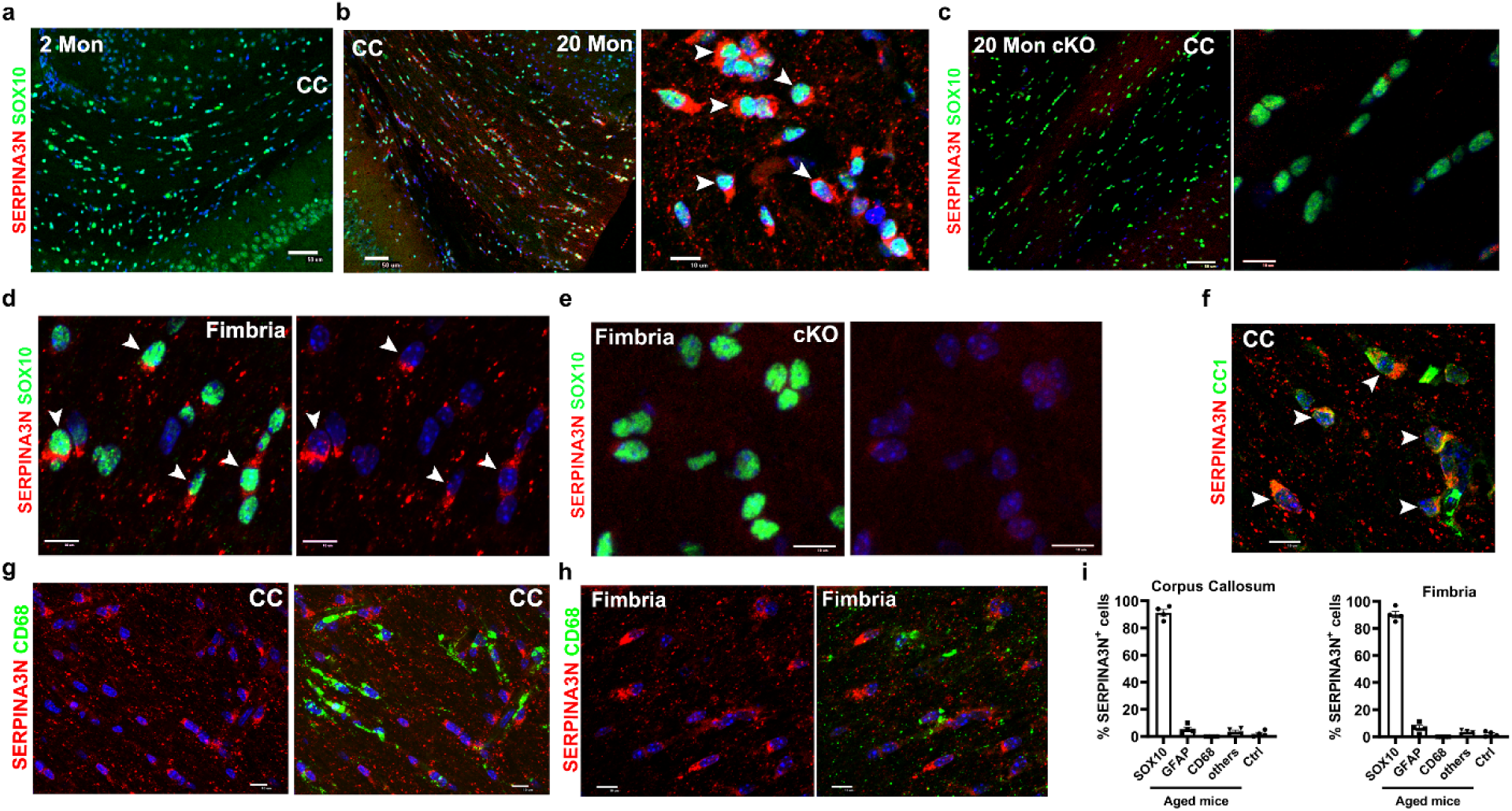
–SerpinOLs are present in the brain of normal aging. **a-c** double IHC of SERPINA3N and SOX10 in the corpus callosum (CC) of young (2 months, Mon, **a**), aged (20 Mon, **b**) and aged mice of *Serpina3n* cKO (**c**). Arrowheads indicate SERPINA3N^+^SOX10^+^ cells. Numerous SERPINA3N puncta were observed in aged mice and abolished in aged *Serpina3n* cKO mice. **d-e** SERPINA3N^+^SOX10^+^ SerpinOLs (arrowheads) in the fimbria of aged WT (**d**) and *Serpina3n* cKO (**e**) mice. **f** Confocal image showing SERPINA3N^+^ cells co-labeled with CC1^+^ mature OLs in aged CC. **g-h** Fluorescent IHC showing absence of SERPINA3N from CD68^+^ cells in aged CC (**g**) and fimbria (**h**). **i** Percentage of SERPINA3N^+^ cells positive for indicated markers in aged and young (Ctl) mice. *n* = 4 mice each group. Data were presented as mean ± s.e.m. Scale bars: **a** 50 µm, **d-h** 20 µm.

We generated transgenic *Olig2-Cre:Serpina3n*^fl/fl^ mice (oligodendroglial *Serpina3n* cKO) to study if OLs are the cellular source of SERPINA3N during normal aging. We found that SERPINA3N immunoreactive signal, both intracellular and extracellular, was largely abolished in the corpus callosum (**Fig. 4c**) and fimbria (**Fig. 4e**) of aged *Serpina3n* cKO mice compared with controls, validating OLs as the primary producers of SERPINA3N in the aged brain.

Normal aging is associated with chronic activation of microglia ^43^. We found that SerpinOLs were markedly increased in the regions of corpus callosum (**Fig. 4g**) and fimbria (**Fig. 4h**) where CD68^+^ activated microglia were frequently observed. The proximity of SerpinOLs with CD68^+^ activated microglia indicates that OL-derived SERPINA3N may regulate microglial behaviors. Collectively, our results establish that normal aging transitions homeostatic OLs into SerpinOLs.

### Neuroinflammation is insufficient to transition homeostatic OLs into SerpinOLs

The mechanisms underlying the transition of homeostatic OLs into SerpinOLs remain poorly understood. It has been proposed that CNS inflammatory environment mediates that transition ^24^. Given that neuroinflammation is a shared hallmark of various neurological conditions, including ischemic stroke, endotoxin exposure, Alzheimer’s disease, neurotrauma, and normal aging, it is a plausible candidate mechanism. To investigate the role of CNS inflammation in this transition, we employed two independent experimental approaches.

In the first approach, mice were inoculated with complete Freund’s adjuvant (CFA), a potent stimulator of the innate immune system ^44^, to elicit CNS inflammation and glial activation. Unbiased RNA-seq analysis (**Fig. 5q**) revealed robust inflammatory responses in the spinal cord of CFA-treated mice, as evidenced by the induction of a large cohort of DEGs (**Fig. 5b, Suppl Table 4**) that participated in immune system processes, innate immune responses, and bacterial responses (**Fig. 5c**). CFA also triggered astrocyte activation, as shown by the upregulation of reactive astrocyte markers (**Fig. 5d**). Despite these marked inflammatory and glial responses, *Serpina3n* transcripts were not induced in the spinal cord of CFA-treated mice compared with PBS controls (**Fig. 5e, f**). We found that CFA treatment did not cause oligodendroglia damage or loss (**Fig. 5g**). These data suggest that neuroinflammation, in the absence of OL injury, is insufficient to transition homeostatic OLs into SerpinOLs.

**Figure 5.**
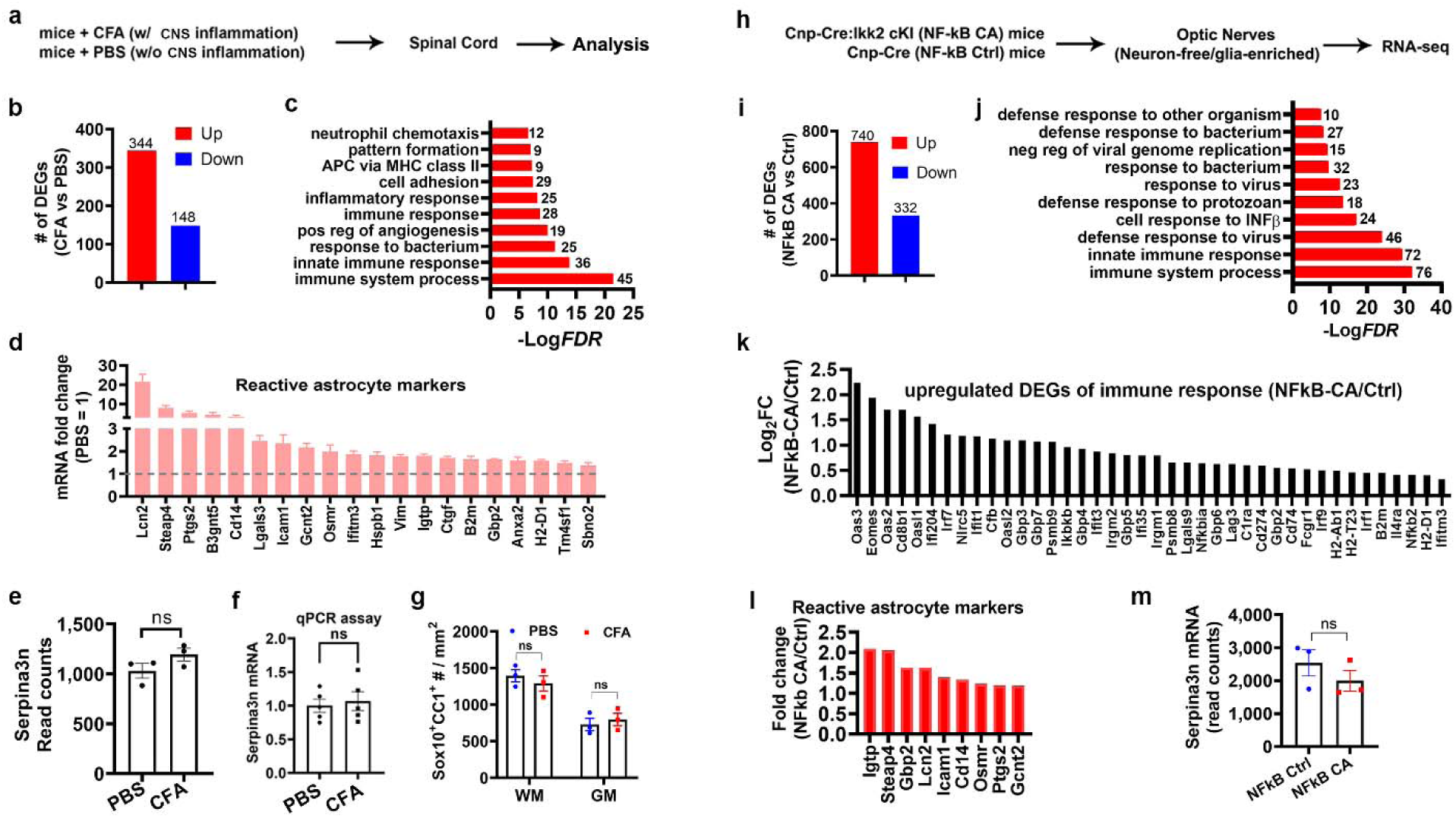
CNS inflammation is not sufficient but promotes SERPINA3N expression in OLs. **a** Experimental design for panels **b-g**. Adult mice were immunized with CFA or PBS (s.c. day 0) and pertussis (i.p. days 0 and 2). Spinal cords were analyzed at day 35 post-CFA (or PBS). **b** Number of up- or -downregulated DEGs in CFA vs. PBS spinal cord identified by RNA-seq (see Suppl Table 4). **c** Top 10 enriched gen ontology (GO) terms overrepresented by upregulated DEGs in CFA vs. PBS (Suppl Table 4). **d** Upregulation of reactive astrocyte marker genes in CFA vs. PBS. **e** *Serpina3n* RNA-seq read counts. *P* = 0.1674. *n* = 3 mice each group. **f** RT-qPCR of *Serpina3n* mRNA. *P* = 0.7005. *n* = 5 mice each group. **g** Densities of OLs (SOX10^+^CC1^+^) in spinal cord white matter (WM) and gray matter (GM). *P* = 0.4753 WM, *P* = 0.6081 GM. *n* = 3 mice each group. **h** Experimental design for panels **i-m**. RNA-seq was performed on optic nerves from oligodendroglial-specific NFkB constitutive activation (NFkB CA) and Ctrl mice. **i** Number of DEGs in NFkB CA vs. Ctrl mice identified by RNA-seq (Suppl Table 5). **j** Top 10 enriched GO terms overrepresented by upregulated DEGs in NFkb CA (Suppl Table 5). **k** Examples of immune response genes upregulated in NFkB CA (Suppl Table 5). **l** Upregulation of reactive astrocyte marker genes in NFkB CA vs. Ctrl mice. **m** *Serpina3n* RNA-seq read counts. *P* = 0.3459. *n* = 3 mice each group. Data were presented as mean ± s.e.m. (e, f, g, m).

In the second approach, we used a genetic model to induce CNS inflammation. We generated *Cnp-Cre:R26Stop^FL^ikk2ca* transgenic mice (referred to as NFkB CA mice) in which NFkB signaling, a key regulator of inflammation^45^, is constitutively activated in oligodendroglial lineage cells ^46^. Our previous studies showed that NFkB activation does not affect oligodendroglial viability under normal conditions ^46^. Our analysis of glial cell-enriched optic nerves (**Fig. 5h**) revealed marked transcriptomic changes (**Fig. 5i, Suppl Table 5**), including upregulation of gene associated with immune system process and innate immune response (**Fig. 5j, k**), astroglial activation (**Fig. 5l**), and many pro-inflammatory mediators (such as CXCL10, CXCL9, CCL3, CCL12, **Suppl Table 5**), compared with NFkB Ctrl mice. Despite this robust inflammatory environment, Serpin3n was not induced in NFkB CA mice (**Fig. 5m**), further supporting the conclusion that CNS inflammation alone does not drive the transition of homeostatic OLs into SerpinOLs. These results suggest that additional factors, such as direct OL injury or damage, play crucial roles in this transition.

### Oligodendroglial injury drives the transition of homeostatic OLs into SerpinOLs

To determine whether OL injury is sufficient to trigger the transition of homeostatic OLS into SerpinOLs, we employed the CPZ demyelinating model during the very acute phase. CPZ diet-mediated demyelination is considered as a primary demyelinating model in which the copper-chelating CPZ diet directly causes OL damage/injury presumably through mitochondrial dysfunction and glial activation is secondary to OL damage/injury (inside-out model) ^47^ particularly during acute phases when glial activation is minimal. Previous studies have shown that CPZ consumption induces rapid OL injury and loss in the corpus callosum as early as 24 hours ^48^. We observed a significant increase in SERPINA3N expression at day 2 of CPZ consumption in the brain (**Fig. 6a, b**). At this early time-point, the activation of microglia or astroglia was yet to be detected, as evidenced by unaltered levels of microglial markers (IBA1 or CD68) and astroglial markers (GFAP) at both protein and histological levels (**Fig. 6a-e**). Notably, SERPINA3N expression was readily detected in SOX10^+^ cells exhibiting “beads-on-a-string” morphology (**Fig. 6f**, boxed area), characteristics of normal OLs in the corpus callosum under homeostatic conditions. To corroborate these findings, we utilized sensitive *Serpina3n*-tdTom reporter mice. tdTom was colocalized with endogenous SERPINA3N at day 2 post-CPZ diet (**Fig. 6g**), confirming the accuracy of the reporter in marking SERPINA3N-producing cells. We found that approximately 20% of SOX10^+^ oligodendrocytes were transitioned into tdTom^+^ SerpinOLs in the corpus callosum at this stage (**Fig. 6h**). Consistent with our findings in 4-week CPZ (**Fig. 1q**, **Fig. 2g**), few tdTom signals were was detected in GFAP^+^ astrocytes at 2 days CPZ (**Fig. 6i, j**). These results suggest that oligodendroglial injury alone is sufficient to drive the transition of homeostatic OLs into SerpinOLs.

**Figure 6.**
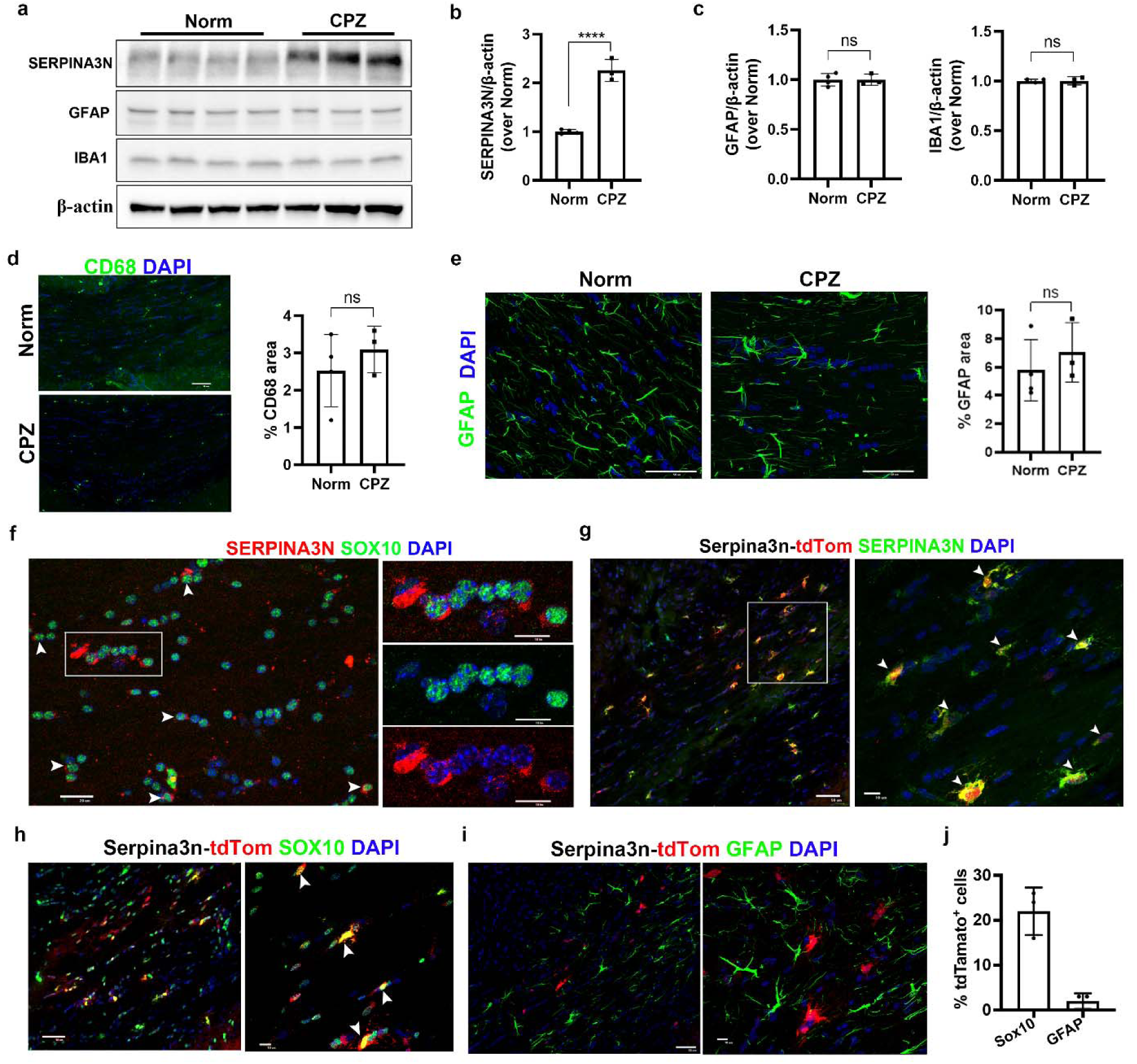
CPZ-induced OL injury/damage triggers transition to SerpinOLs. **a-c** Western blot and quantification of SERPINA3N, GFAP, and IBA1 in brains of adult mice after 2 days of CPZ diet. **** *P* < 0.0001 SERPINA3N; *P* = 0.9916 GFAP; *P* = 0.9097 IBA1. *N* = 4 mice for Norm; *n* = 3 mice for CPZ. **d-e** IHC and quantification of CD68^+^ (**d**) and GFAP^+^ cells I in the corpus callosum after 2 days CPZ. *P* =0.4172 CD68; *P* = 0.4748 GFAP. *N* = 4 mice for Norm; *n* = 3 mice for CPZ. **f** IHC of SERPINA3N and SOX10 in the corpus callosum after 2 days of CPZ. Arrowheads indicate SERPINA3N^+^ OLs. Boxed region shown at higher magnification. g IHC of SERPINA3N and tdTom in the corpus callosum of Serpina3n-tdTom mice on CPZ diet for 2 days. Boxed region shown at higher magnification. Arrowheads point to double positive cells. **h-i** IHC of tdTom with SOX10 (**h**) or GFAP(**i**) in the corpus callosum of Serpina3n-tdTom mice on CPZ diet for 2 days. **j** Quantification of the percentage of tdTomato^+^ cells co-expressing SOX10 or GFAP after 2 days of CPZ. *N* = 3 mice for CPZ. Scale bars: **d-e** 50 µm, **f** 20 µm, **g-I** 50 µm (lower magnification) and 10 µm (higher magnification). Data were presented as mean ± s.d. (b, c, d, e, j)

Collectively, our findings suggest that oligodendroglial injury, rather than neuroinflammation or glial activation, may be a possible direct mechanism underlying the transition of homeostatic OLs into SerpinOLs.

### SerpinOLs exhibit inflammation/immune regulatory signatures and STAT3 signaling activation

The molecular signatures of SerpinOLs and underlying molecular mechanisms of SerpinOL transition remain incompletely understood. To investigate these, we performed single cell RNA sequencing (scRNA-seq) on spinal cord cells from MOG/EAE mice to characterize SerpinOLs under demyelinating conditions. From 7,946 high-quality live cells pooled from three spinal cords, we identified 49 distinct clusters (C0–C48) based on cluster-specific marker genes (**Fig. 7a, Suppl Table 6**). Seven clusters (C6, C0, C2, C14, C33, C39, and C18) were annotated as oligodendroglial lineage cells (**Fig. 7b, Suppl Fig. 10**). Consistent with previous findings, *Serpina3n* expression was largely confined to oligodendrocyte clusters (C6, C0, C2, C14, C33) (**Fig. 7c**). We defined SerpinOLs and non-SerpinOLs based on their Serpina3n expression and conducted pseudo-bulk differential expression analysis. This revealed 126 DEGs in SerpinOLs compared with non-SerpinOLs under MOG/EAE conditions (**Fig. 7d**, **Suppl Table 7**). Functional annotation of these DEGs showed significant enrichment for pathways related to immune response, immune regulation, JAK-STAT signaling (**Fig. 7e, Suppl Table 7**), a pathway critically involved in inflammation modulation ^49^. Notable DEGs included those with established roles in immune regulation and activation such as *Apod* ^50^, *Rb1* ^51^, *Nfat5* ^52,53^, *Klk6* ^54^, and *Sbno2* ^55–57^ (**Fig. 7d**). While prior studies have shown that OL lineage cells acquire immune cell-like phenotypes including MHC-II molecule expression in MOG/EAE ^58,59^, We found that few SerpinOLs express immune cell markers such CD45 (**Fig. 7f**) or key MHC-II molecules such as I-A/I-E (**Fig. 7g**) and CD74 (**Fig. 7h**). These findings suggest that SerpinOLs are molecularly distinct from previously reported populations of antigen phagocytosing and presenting properties ^59^ and may exert immune-regulatory functions through MHC-II-independent mechanisms.

**Figure 7.**
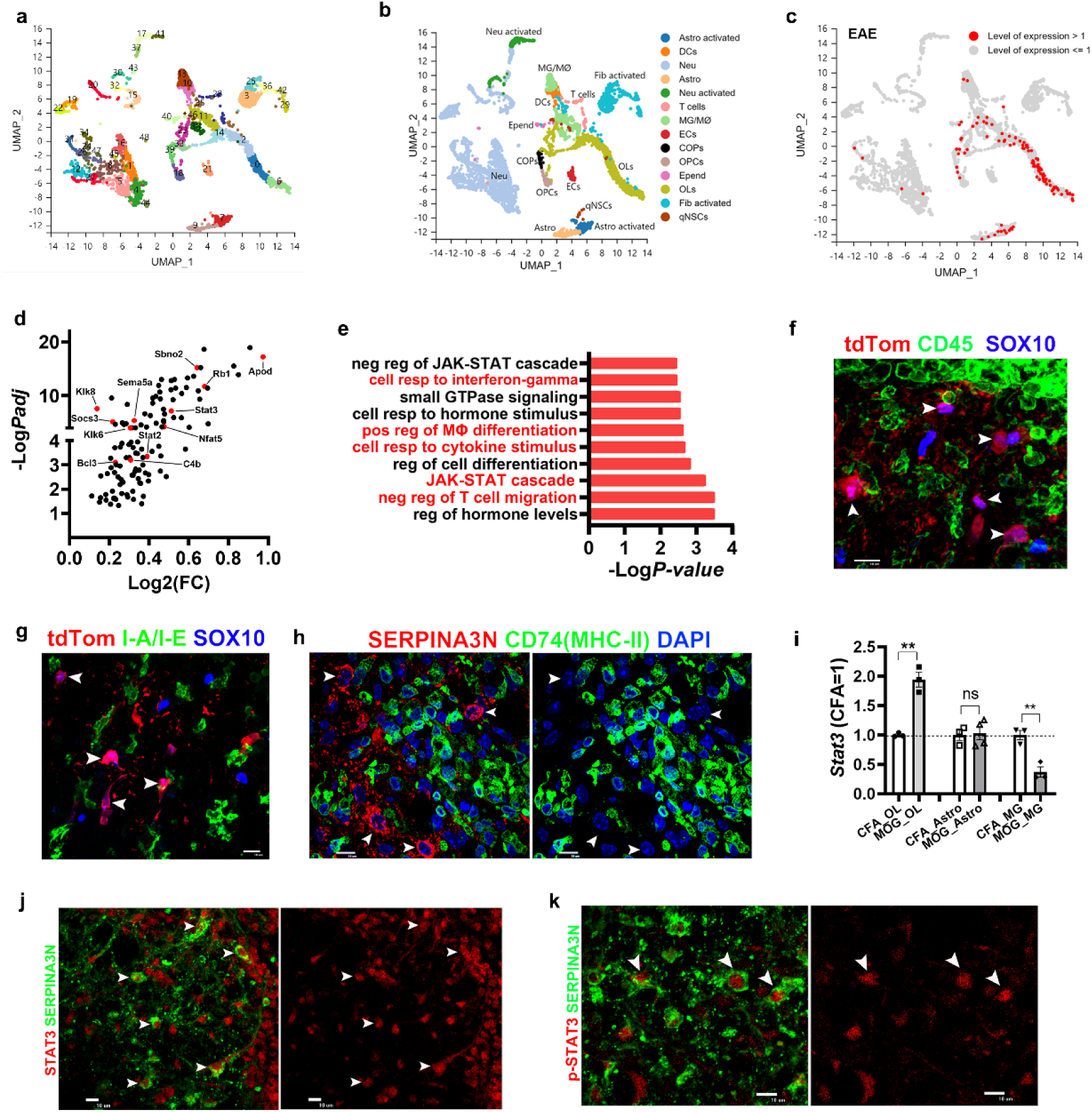
molecular characterization of SerpinOLs in demyelinating injury. **a-b,** UMAP clusters (**a,** Suppl Table 6) and cell type annotations (**b**) of 7,946 single cells (scRNA-seq) from the spinal cord at D30 post-MOG/EAE (pooled three spinal cords). MG/MØ, microglia/macrophages, DCs, dendritic cells; Neu, neurons; Astro, astrocytes; ECs, endothelial cells; Epend, ependymal cells; OPCs, oligodendrocyte progenitor cells; COPs, differentiation-committed OPCs; OLs, oligodendrocytes; Fib, fibroblasts; qNSC, quiescent neural stem cells. **c**, *Serpina3n* expression levels across different cell populations. *Serpina3n* mRNA is highly enriched in the OL clusters. **d**, significantly upregulated genes in SerpinOLs compared with non-Serpina3n-expressing OLs, assessed by pseudobulk RNA analysis (Suppl Table 7). **e**, Example of biological processes overrepresented by upregulated genes in SerpinOLs (Suppl Table 7). **f,** triple IHC showing that tdTom^+^SOX10^+^ SerpinOLs (arrowheads) are negative for CD45 in MOG/EAE-treated Serpina3n-tdTom reporter mice at day 21. Scale bar=10µm. **g,** triple IHC showing that tdTom^+^SOX10^+^ SerpinOLs (arrowheads) do not express I-A/I-E in MOG/EAE-treated Serpina3n-tdTom reporter mice at day 21. Scale bar=10µm. **h,** confocal images showing that SERPINA3N^+^ cells (arrowheads) are negative for MHC-II molecule CD74 at day 21 post-MOG/EAE. Scale bar=10µm. **i**, normalized read counts of Stat3 in RNA-seq of purified glial cell populations in MOG/EAE and CFA mice at D30. n = 3 each population except MOG_Astro (n = 4), Error bars, s.e.m. **j-k**, colocalization of SERPINA3N with STAT3 (**j**, arrowheads) and active phosho-STAT3 (**k**, arrowheads) in D30 MOG/EAE spinal cord. Scale bar=10µm.

Importantly, we found that JAK/STAT signaling components and target genes (*Stat3, Stat2, Socs3, Bcl3, and Sbno*) were significantly enriched in SerpinOLs (**Fig. 7d, e**). Using unbiased RNA-seq of purified glial populations, we found that Stat3 mRNA was upregulated preferentially in OLs under inflammatory demyelinating conditions (**Fig. 7i**). We used IHC to confirm the colocalization of STAT3 and SERPINA3N. Our results showed that nearly all SERPINA3N^+^ cells expressed STAT3 (**Fig. 7j**). STAT3 is activated by phosphorylation at Tyr705 which induces dimerization, nuclear translocation, and DNA binding ^60^. We then used the signaling active form of STATs, phosphorylated form, p-STAT3 (Tyr705) to characterize SerpinOLs and found that all SERPINA3N-expressing cells were positive for p-STAT3 (**Fig. 7k**) in MOG/EAE spinal cords, suggesting that SerpinOLs exhibit the activity of the STAT3-mediated signaling. Our finding is consistent with previous data showing that STAT3 binds to the promoter region of SERPINA3 gene in cancer cells ^61^. Collectively, our findings indicate that SerpinOLs may gain the function of regulating neuroinflammation during diseased conditions and this regulatory role may be independent of MHC-II expression as previously reported. Our results also suggest that STAT3 signaling may be an upstream molecular trigger for SerpinOL transition.

### SerpinOLs regulate CNS inflammation and microglial activation during demyelination

To determine whether SerpinOLs regulate CNS inflammatory responses, we generated transgenic mice with oligodendrocyte-specific SERPINA3N deficiency and subjected them to MOG/EAE demyelinating injury. scRNA-seq analysis (**Fig. 8a**) revealed a significant increase in the proportion of myeloid cells (**Fig. 8b**), including DCs (C10) and microglia/macrophages (MG/MØ; C11, C13, and C38) (**Suppl Fig. 11a-c)**, in the spinal cords of MOG versus CFA Ctrl mice, supporting the known role of myeloid cell-mediated CNS inflammation in MOG/EAE demyelination ^28^. Strikingly, SERPINA3N ablation significantly reduced the myeloid cell number in the spinal cords of cKO_MOG mice compared with Ctrl_MOG mice (**Fig. 8c**). These findings were in congruent with scRNA-seq results showing decreased expression of myeloid cell marker CD68 (**Suppl Fig. 11d**), suggesting that SerpinOLs mediate CNS inflammatory response to MOG/EAE injury through producing SERPINA3N.

**Figure 8.**
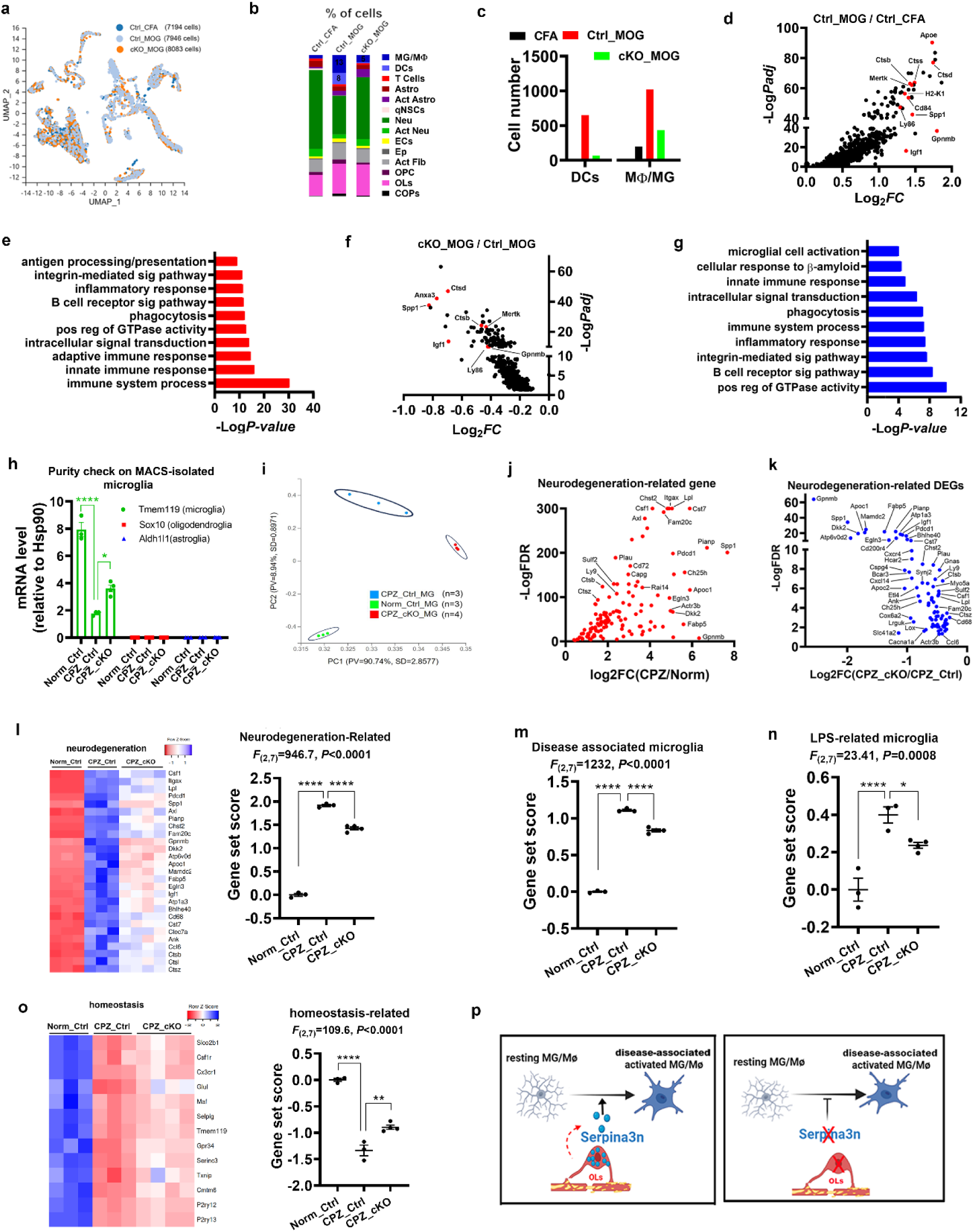
SerpinOLs regulate CNS inflammation and microglial activation in response to demyelination. **a** UMAP clusters of single cells from spinal cord of three groups of mice at D30 post-MOG/EAE (or CFA). **b** Percentage of cell types. **c** Numbers of dendritic cells (DCs) and microglia/macrophages (MG/MØ) in each group. **d** Upregulated DEGs in Ctrl_MOG vs. Ctrl_CFA revealed by psuedobulk analysis of MG/MΦ (C11, C13, C38) (Suppl Table 8). **e** Biological processes of upregulated DEGs in Ctrl_MOG mice (Suppl Table 8). **f** Downregulated DEGs in cKO_MOG versus Ctrl_MOG mice revealed by psuedobulk analysis of MG/MΦ (Suppl Table 9). g Biological processes of downregulated DEGs in Ctrl_MOG mice (Suppl Table 9). **h** purity check of microglia (MG) by RT-qPCR assay of lineage-specific marker genes. MG were purified from *Serpina3n* Ctrl brain of normal diet (Norm_Ctrl, n = 3) and 4 weeks CPZ (CPZ_Ctrl, n = 3) and *Serpina3n* cKO brain of 4 weeks CPZ (CPZ_cKO, n = 4)). Microglial RNA was used for RNA-seq and bioinformatics in Panels **k-q**. **** *P* < 0.0001, * *P* = 0.0104. **i** Principal component analysis of MG samples. N = 3 mice Norm_Ctrl and CPZ_Ctrl; *n* = 4 mice CPZ_cKO **j** Upregulation of core neurodegeneration-related genes^62^ in MG of CPZ_Ctrl versus Norm_Ctrl mice (Suppl Table 10). k Downregulation of core neurodegeneration-related genes^62^ in MG of CPZ_cKO versus CPZ_Ctrl mice (Suppl Table 10). **l,** heatmap and cumulative scores of core neurodegeneration-related gene set (Suppl Table 11). One way ANOVA and Tukey’s post-hoc test, *P* < 0.0001 CPZ_Ctrl vs Norm_Ctrl, *P* < 0.0001 CPZ_cKO vs CPZ_Ctrl. N = 3 mice Norm_Ctrl and CPZ_Ctrl; *n* = 4 mice CPZ_cKO **m-n,** cumulative scores of disease-associated (**m**) ^77^ and LPS-related (**n**) ^62^ microglial gene sets (Suppl Table 11) showing an attenuated transcriptomic response of microglia to demyelination in CPZ_cKO mice. One way ANOVA and Tukey’s post-hoc test, **** *P* < 0.0001, *** *P* = 0.0006, * *P* = 0.0493. N = 3 mice Norm_Ctrl and CPZ_Ctrl; *n* = 4 mice CPZ_cKO **o,** heatmap and cumulative scores of homeostatic microglial gene set (Suppl Table 11). One way ANOVA and Tukey’s post-hoc test, *P* < 0.0001 CPZ_Ctrl vs Norm_Ctrl, *P* = 0.0034 CPZ_cKO vs CPZ_Ctrl. N = 3 mice Norm_Ctrl and CPZ_Ctrl; *n* = 4 mice CPZ_cKO **p,** graphic conclusion of microglial RNA-seq. Oligodendrocyte-derived SERPINA3N promotes the activation of homeostatic microglia towards disease-associated and neurodegeneration-related states. Data were presented as mean ± s.e.m. (h, i, m, n, o).

To further explore this mechanism, we conducted pseudo-bulk RNA-seq analysis of MG/MØ clusters (C11, C13, C38). In Ctrl_MOG mice, 947 genes were significantly upregulated compared with Ctrl_CFA mice (**Fig. 8d**, **Suppl Table 8**), which are primarily related to inflammation, immune response, and phagocytosis (**Fig. 8e**). Interestingly, of the 947 DEGs, 417 (44%) were significantly downregulated in MG/MØ of cKO_MOG versus Ctrl_MOG mice (**Fig. 8f**), representing 79% of the 530 total downregulated DEGs (417/530) in the cKO_MOG condition (**Suppl Table 9**). These 417 genes were primarily associated with inflammatory/immune responses and phagocytosis among others (**Fig. 8g**), indicating that SERPINA3N deficiency dampens the pro-inflammatory transcriptomic profile of MG/MØ in MOG/EAE. Given that myeloid cells do not express SERPINA3N, our findings suggest that SERPINA3N secreted from SerpinOLs may act in a paracrine manner to promote MG/MØ-driven neuroinflammation during EAE demyelinating injury.

To further validate the role of SerpinOLs in neuroinflammation and microglial activation, we employed CPZ demyelinating model in which microglia, but not peripheral monocytes, drive neuroinflammation (cf **Suppl Fig. 3a**). RNA-seq was performed on microglia purified from brains of healthy controls (Norm_Ctrl), CPZ-treated controls (CPZ_Ctrl), and CPZ-treated Serpina3n cKO (CPZ_cKO) mice (**Fig. 8h**). PCA analysis showed that SERPINA3N deletion markedly shifted the microglial transcriptomic profiles away from that of the CPZ_Ctrl group (**Fig. 8i**), indicating an altered activation state.

A previously defined microglial gene module comprising 134 core neurodegeneration-related genes ^62^ was used to assess disease-related microglial activity. Of these, 125 (93%) were upregulated in microglia of CPZ_Ctrl versus Norm_Ctrl groups (**Fig. 8j**, **Suppl Table 10**), which is consistent with the finding that activated microglia are essential for CPZ-mediated demyelination ^63^. Remarkably, 81 of these 125 genes (64.8%) were significantly downregulated in CPZ_cKO versus CPZ_Ctrl mice (**Fig. 8k, Suppl Table 10**), suggesting that SERPINA3N deficiency attenuates neurodegeneration-associated microglial activation. To further characterize microglial state, we calculated cumulative gene set scores (**Suppl Table 11;** see Materials and Methods). Gene set associated with neurodegeneration (**Fig. 8l**), disease-associated microglia (**Fig. 8m**), and LPS-induced proinflammatory activation (**Fig. 8n**) were all significantly decreased in CPZ_cKO versus CPZ_Ctrl microglia. In contrast, the homeostasis-related gene set score was significantly increased in CPZ_cKO microglia (**Fig. 8o**). These data support a working model in which SERPINA3N depletion in SerpinOLs shifts microglia from a disease-associated and proinflammatory state toward a more homeostatic state in response to demyelination (**Fig. 8p**). Collectively, our findings establish that injury-transduced SerpinOLs regulate CNS inflammation and promote microglial activation toward disease-associated and neurodegeneration-related activation state via SERPINA3N during CNS demyelination.

### SerpinOLs regulate neuroinflammation and glial activation in non-demyelinating conditions

We next investigated whether the role of SerpinOLs in regulating neuroinflammation is preserved in non-demyelinating conditions. Given our findings that homeostatic OLs transition into SerpinOLs during normal aging, we hypothesize that SerpinOLs contribute to age -related neuroinflammation and glial activation. To test this hypothesis, we examined the brains of young (2 mon), aged (20 mon), and aged Serpina3n cKO (20 mon, *Olig2-Cre:Serpina3n*^fl/fl^) mice. Oligodendroglia-specific deletion of SERPINA3N was confirmed by Western blot (**Fig. 9a, b**) and histological (cf **Fig. 4c, e**) assays. Notably, depleting SERPINA3N attenuated aging-elicited activation of microglia and astrocytes, as evidenced by reduced expression of microglial CD68 and IBA1 and astroglial GFAP at both mRNA and protein levels in aged cKO brains compared with age-matched Ctrl mice (**Fig. 9a-c**). Histological quantification confirmed that SERPINA3N deletion normalized CD68^+^ area in the corpus callosum of aged cKO mice to levels observed in young mice (**Fig. 9d**). Aging is known to drive astrocyte activation toward the neurotoxic A1 phenotype ^38^; this was supported by increased expression of the representative pan-reactive astrocytes marker *Cxcl10* and neurotoxic reactive astrocyte marker *C3* (**Fig. 9e**). Impressively, SERPINA3N deficiency significantly reduced the expression of these markers in aged cKO brain (**Fig. 9e**). Additionally, expression of aging-elicited pro-inflammatory cytokine *Il1b* and chemokine *Ccl2* was significantly decreased in aged cKO mice (**Fig. 9f**), alongside reduced expression of the M1-polarized pro-inflammatory microglial marker CD86 (**Fig. 9g**). These results suggest that OL-derived SERPINA3N promotes aging-associated neuroinflammation and glial activation toward pro-inflammatory and neurotoxic states.

**Figure 9.**
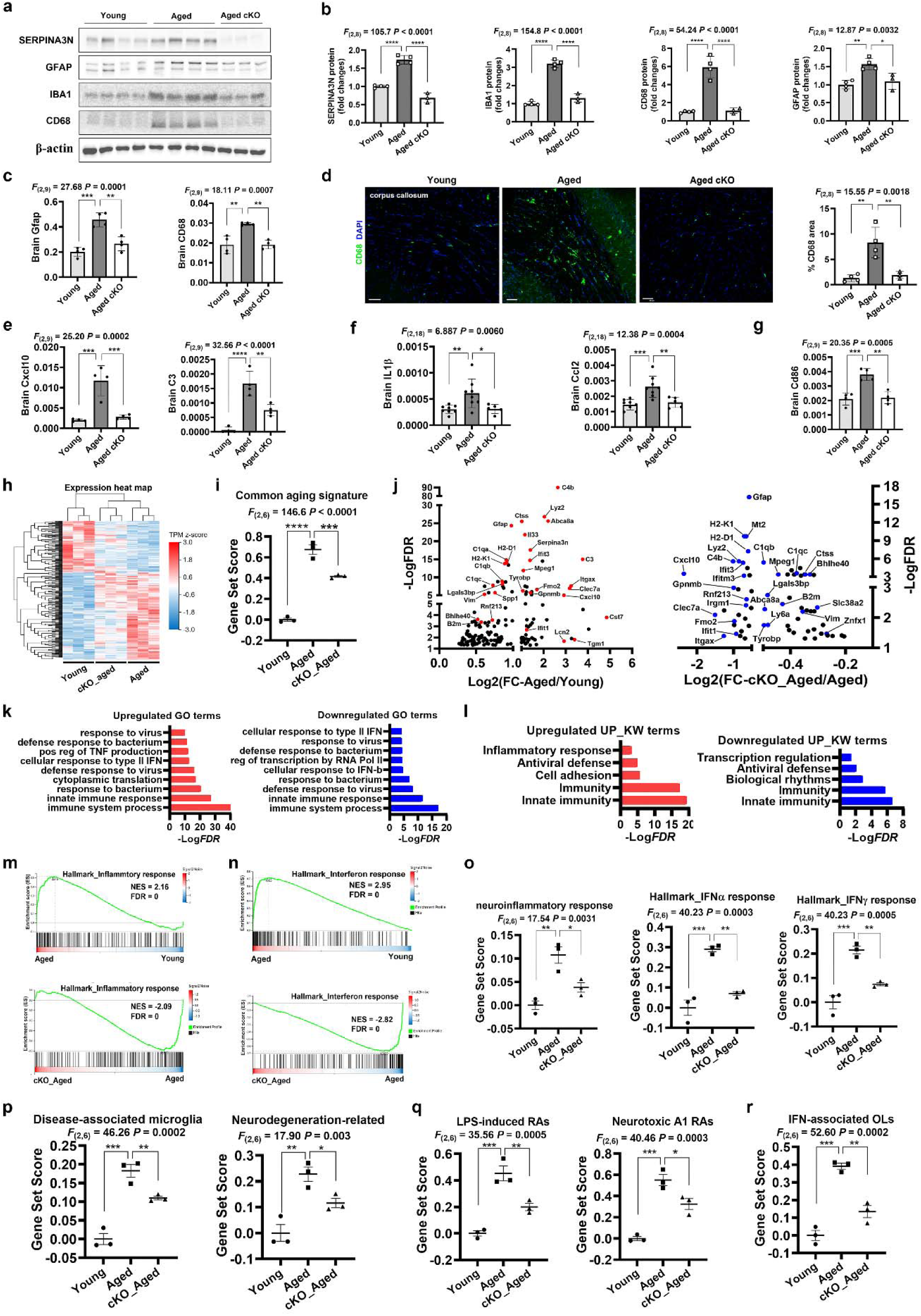
SerpinOLs regulate CNS inflammation and glial responses during normal aging. **a-b**, Western blot and quantification of SERPINA3N, IBA1, CD68, and GFAP in the brains of young (2 mon, n = 4), aged (20 mon, n = 4), and Serpina3n cKO aged (20 mon, n = 3) mice. One way ANOVA followed by Tukey’s multiple comparisons test. SERPINA3N **** *P* < 0.0001, IBA1 *\*\*\*\*P* < 0.0001, CD68 *\*\*\*\* P* < 0.0001, GFAP ** *P* = 0.0035, * *P* = 0.0147. **c**, RT-qPCR of brain *Gfap* and *Cd68* mRNA levels. One way ANOVA followed by Tukey’s multiple comparisons test. *Gfap* *** *P* = 0.001, ** *P* = 0.0012, Cd68 ** *P* = 0.0014. *n* = 4 mice each group **d**, fluorescent IHC and quantification of CD68. Scale bar=50µm. One way ANOVA followed by Tukey’s multiple comparisons test. ** *P* = 0.0023 Aged (n = 4) vs Young (n = 4), ** *P* = 0.0063 cKO_Aged (n = 3) vs Aged (n = 4). **e**, RT-qPCR assay for the brain levels of pan reactive astrocyte marker Cxcl10 and neurotoxic A1 polarized reactive astrocyte marker C3. One way ANOVA followed by Tukey’s multiple comparisons test. Cxcl10 *** *P* = 0.0003 Aged vs Young, *** *P* = 0.0006 cKO_Aged vs Aged; C3, **** *P* < 0.0001, ** *P* = 0.0033. *n* = 4 mice each group **f**, brain mRNA levels of pro-inflammatory mediators Ccl2, and IL1β assessed by RT-qPCR. One way ANOVA followed by Tukey’s multiple comparisons test. Ccl2 *** *P* = 0.0005, ** *P* = 0.006, Il1b ** *P* = 0.0092, * *P* = 0.0257. n = 8 mice Young and Aged; n = 5 mice Aged_cKO. **g**, RT-qPCR assay for the brain levels of pro-inflammatory M1 polarized microglial marker CD86. One way ANOVA followed by Tukey’s multiple comparisons test. *** *P* = 0.0008, ** *P* = 0.0012. *n* = 4 mice each group **h**, unsupervised clustering heatmap of DEGs in the brains of young (2 mon), aged (20 mon), and Serpina3n cKO aged (20 mon) mice (Suppl Table 12). **i**, cumulative scores of common aging signature gene set ^41^ (Suppl Table 11). One way ANOVA followed by Holm-Sidak multiple comparisons test. **** *P* < 0.0001, *** *P* = 0.0006. *n* = 3 mice each group **j**, upregulated DEGs in the aged brain versus young (left) and downregulated DEGs in cKO_aged brain versus aged (right). Most upregulated genes associated with immune/inflammatory response and microglial/astroglial activation in aged (vs young) brain were downregulated in cKO_aged brain. **k**, GO analysis showing biological processes overrepresented by upregulated DEGs in aged versus young (left, red) and by downregulated DEGs in cKO_aged versus aged (right, blue). **l**, uniport keyword biological processes (UP_KW_BP) overrepresented by upregulated DEGs in aged versus young (left, red) and by downregulated DEGs in cKO_aged versus aged (right, blue). **m-n**, gene set enrichment assay (GSEA) of MSigDB Hallmark Pathways showing that inflammatory response (**m**) and interferon response (**n**) were significantly upregulated in aged brain versus young (upper panels) but downregulated in cKO_aged brain versus aged (lower panels). NES, normalized enrichment score. **o,** cumulative gene set scores of neuroinflammatory and interferon responses. One way ANOVA followed by Holm-Sidak multiple comparisons test. Neuroinflammatory response, ** *P* = 0.0033, * *P* = 0.0183; IFNα response, *** *P* = 0.0004, ** *P* = 0.0013; IFNγ response, *** *P* = 0.0005, ** *P* = 0.0034. *n* = 3 mice each group. **p,** cumulative scores of disease-associated microglia and neurodegeneration-related microglia gene sets. One way ANOVA followed by Holm-Sidak multiple comparisons test. DAM *** *P* = 0.0002, ** *P* = 0.0088; NDM ** *P* = 0.0022, * *P* = 0.0451. *n* = 3 mice each group. **q,** gene set scores of LPS-induced reactive astrocytes ^35^ and neurotoxic A1 reactive astrocytes ^78^ (Suppl Table 11). One way ANOVA followed by Holm-Sidak multiple comparisons test. LPS-induced RAs *** *P* = 0.0005, ** *P* = 0.0066, A1 RAs *** *P* = 0.0003, * *P* = 0.0105. *n* = 3 mice each group. **r,** gene set scores of interferon-responsive OLs in each group of mice. One way ANOVA followed by Holm-Sidak multiple comparisons test. IFN *** *P* = 0.0002, ** *P* = 0.0011, DA1 *P* = 0.6777 aged vs young, *P* = 0.8288 cKO_aged vs aged. *n* = 3 mice each group. Data presentation: mean ± s.d. (b, c, d, e, f, g); mean ± s.e.m. (o, p, q, r)

We next performed unbiased brain RNA-seq to strengthen our conclusions. Unsupervised heatmap clustering of DEGs (**Suppl Table 12**) revealed distinct separation between aged cKO and aged Ctrl mice (**Fig. 9h**), suggesting that SERPINA3N deletion alters brain transcriptomics. We examined the expression of a recently identified common aging signature (CAS) consisting of 82 differentially regulated genes in the murine brain ^41^ and found that SERPINA3N deletion significantly reversed this common aging signature (**Fig. 9i**). Specifically, most DEGs induced by normal aging (**Fig. 9j**, left) were significantly downregulated in Serpina3n cKO mice (**Fig. 9j**, right), including disease-associated/neurodegeneration-related microglial signature genes (*Spp1, Ctss, Itgax, Clec7a, Gpnmb, Bhlhe40*, *Lyz2, Tyrobp, and Cts7*), complement genes (*C1qa, C1qb, C1qc, C4b*, and *C3*), pan-reactive astrocyte marker genes (*Lcn2, Gfap,* and *Vim*), and A1 neurotoxic reactive astrocyte marker genes (*C3, B2m, Ifit3, Ifit1, H2-D1,* and *H2-K1*) (**Fig. 9j**). GO analysis revealed that normal aging activated innate immune response and inflammatory response (**Fig. 9k-l**, red), whereas SERPINA3N deficiency significantly reduced these pathways in aged cKO brain (**Fig.9k-l**, blue). These findings indicate that SERPINA3N accelerates brain aging through promoting CNS inflammation and glial activation.

To further corroborate our findings, we performed gene set enrichment assay (GSEA) using the MsigDB molecular signature database. Our results showed that SERPINA3N depletion reversed brain inflammatory response (**Fig. 9m**) and interferon response (**Fig. 9n**) in Serpina3n cKO mice compared with aged mice, as confirmed by significantly rescued gene set scores of neuroinflammatory and interferon type I (IFNα) and type II (IFNγ) responses (**Fig. 9o**). These data suggest that SERPINA3N promotes aging-associated neuroinflammation and cellular responses to interferons, crucial cytokines for innate immune activation.

Microglial and astroglial activation are key mediators of age-related neuroinflammation ^64,65^. We found that SERPINA3N deficiency dysregulated microglial cell activation (**Suppl Fig. 12a**) and significantly decreased the activation states of disease-associated and neurodegeneration-related microglia (**Fig. 9p**), as well as LPS-induced pro-inflammatory response (**Suppl Fig. 12b**) and interferon-related response (**Suppl Fig. 12c**) with minimal effect on microglial proliferation (**Suppl Fig. 12d**) and homeostasis (**Suppl Fig. 12e**). Similarly, SERPINA3N deletion mitigated aging-elicited astrocyte reactivity, including LPS-induced pro-inflammatory response ^35^ and A1 neurotoxic phenotypes ^38^ (**Fig. 9q**). These findings indicate that SerpinOL-derived SERPINA3N is a key regulator of aging-elicited glial activation.

Previous studies have reported that OLs are activated into different disease-associated subpopulations with distinct transcriptomic signatures, including DA1 OLs (immunogenic), DA2 OLs (survival/differentiation) and interferon-responsive OLs ^18,22^. We found that normal aging predominantly induced an IFN-responsive OL state (**Fig. 9r**) in the brain, with no significant enrichment of DA1 or DA2 signatures (**Suppl Fig. 12f-g**). Importantly, depleting SERPINA3N significantly reduced the activation state of interferon responsive OLs (**Fig. 9r**), suggesting that SERPINA3N modulates oligodendroglial responses to interferon during normal aging ^22^. Taken together, our data demonstrate that SerpinOLs, acting through SERPINA3N production, regulate neuroinflammation and promote glial activation toward neurodegenerative, neurotoxic, and interferon-responsive states during normal aging.

### SERPINA3N promotes microglial activation toward pro-inflammatory states

To determine whether SERPINA3N directly promotes pro-inflammatory microglial activation, we utilized primary microglial cultures and a SERPINA3N gain-of-function approach. RT-qPCR assay confirmed the isolated primary microglia were free of astrocyte contamination (**Fig. 10a**) and did not express SERPINA3N under unstimulated (PBS), LPS-stimulated, and SERPINA3N-treated conditions (**Fig. 10b**). As expected, LPS stimulation significantly upregulated CD68, a canonical microglial activation marker while reducing P2ry12, a marker of homeostatic microglia (**Fig. 10c**). We found that treatment with recombinant SERPINA3N further enhanced CD68 expression (**Fig. 10c**), suggesting that SERPINA3N potentiates microglial activation. In addition, SERPINA3N significantly upregulated pro-inflammatory cytokines TNFα and IL-1β (**Fig. 10d**), as well as pro-inflammatory markers iNOS and CD86 (**Fig. 10e**). In contrast, it suppressed the expression of anti-inflammatory markers CD206 and Ym1 (**Fig. 10f**), consistent with a shift toward a pro-inflammatory phenotype. Microglial senescence is increasingly recognized as a pathological feature of the aged brain ^66,67^. We found that SERPINA3N treatment markedly increased the expression of p21 (*Cdkn1a*) (**Fig. 10g**), a well-established marker of cell senescence ^68^ in LPS-stimulated microglia. Collectively, these findings demonstrate a direct effect of SERPINA3N on microglial activation and suggest that SERPINA3N promotes microglial activation toward a pro-inflammatory state.

**Figure 10.**
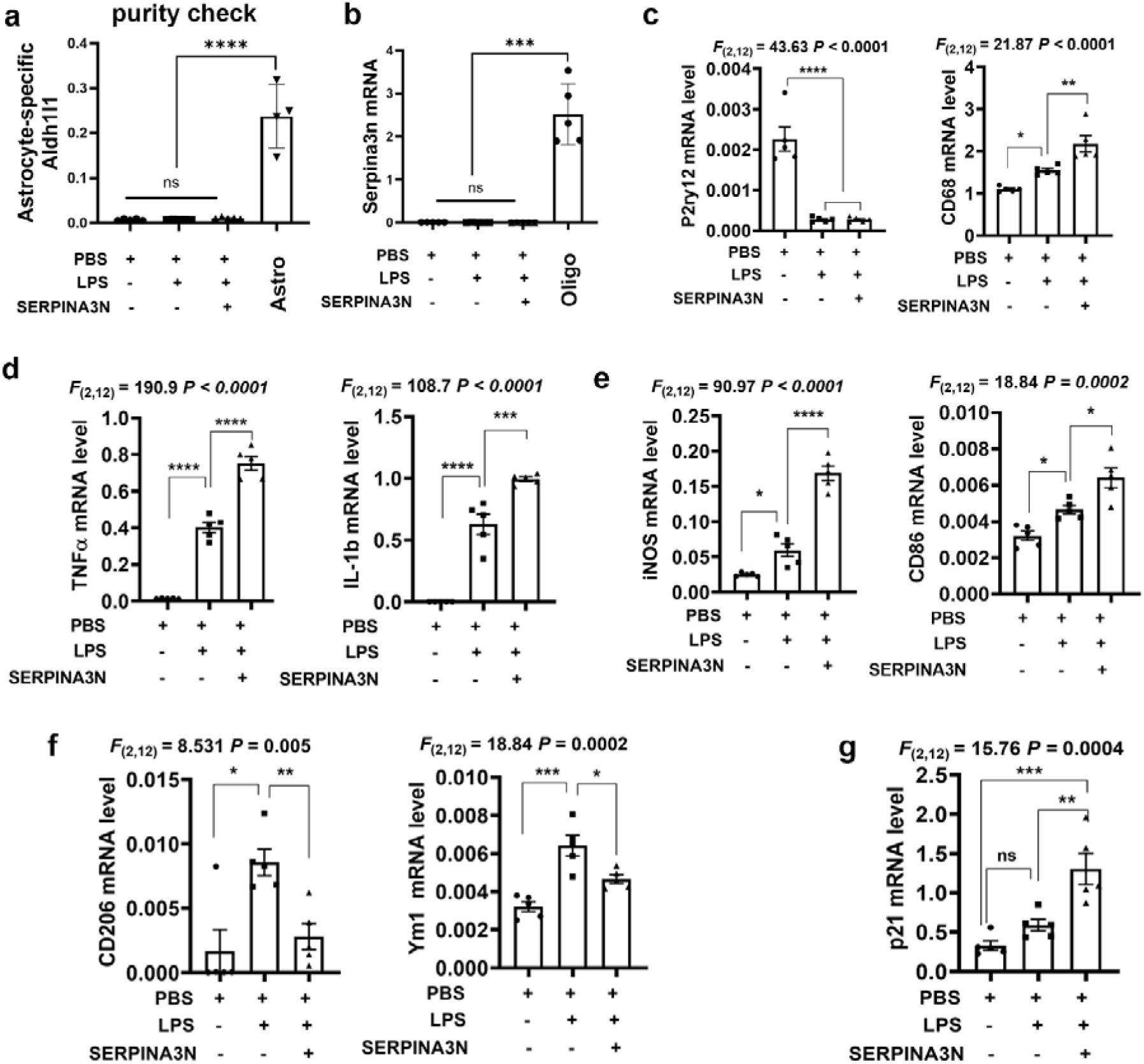
SERPINA3N promotes activated microglia toward a pro-inflammatory state. Primary microglia were isolated from P2 neonatal mice and treated with LPS (10 ng/mL) +/- recombinant mouse SERPINA3N (#4709-PI-010, R&D Systems) at 50ng/mL in culture media for 24 hours prior to RNA isolation (n = 5 independent replicates). **a**, purity assessment by RT-qPCR of astrocytic marker gene *Aldh1l1*. Primary astrocytes (Astro, n = 4 independent replicates) were used as a positive control for Aldh1l1 detection. Mean ± sd. One way ANOVA followed by Tukey’s post-test, *F*_(3, 15)_=54.98, **** *P* < 0.0001 **b**, absence of *Serpina3n* mRNA in cultured microglia, assessed by RT-qPCR. Primary oligodendrocytes (Oligo, n = 5 independent replicates) were used as a positive control for Serpina3n expression. Mean ± sd. One way ANOVA followed by Tukey’s post-test, *F*_(3, 16)_=63.42, *** *P* < 0.001. **c**, RT-qPCR of CD68 and homeostatic marker P2ry12. One way ANOVA followed by Tukey’s test for multiple comparisons. P2ry12 **** *P* < 0.0001; CD68 * *P* = 0.046 LPS vs PBS, * *P* = 0.0059 LPS vs LPS+SERPINA3N **d**, RT-qPCR of canonical pro-inflammatory cytokines TNFα and IL-1β. One way ANOVA followed by Tukey’s post-test. One way ANOVA followed by Tukey’s test for multiple comparisons. TNFα **** *P* < 0.0001; IL-1b *** *P* = 0.0005, **** *P* < 0.0001. **e-f**, RT-qPCR of pro-inflammatory microglial markers iNOS and CD86 (**E**) and anti-inflammatory microglial markers CD206 and Ym1 (**F**). One way ANOVA followed by Tukey’s test for multiple comparisons, iNOS * *P* = 0.0228, **** *P* < 0.0001; CD86 * *P* = 0.0428 LPS vs PBS, * *P* = 0.0142 LPS vs LPS+SERPINA3N; Cd206 * *P* = 0.0188, ** *P* = 0.0060; Ym1 *** *P* = 0.0001, * *P* = 0.0142 **g**, RT-qPCR for the cell senescence marker p21 (CDKN1A). One way ANOVA followed by Tukey’s test for multiple comparisons, ns *P* = 0.3568, ** *P* = 0.0047, *** *P* = 0.0004. Data were presented as mean ± s.d.

## Discussion

Oligodendroglial lineage cells (OPCs and OLs) transition into distinct states identified by their gene expression profiles in response to CNS pathologies. Recently, a subpopulation of immune oligodendroglial lineage cells has been proposed in demyelinating diseases and animal models ^20,58,69,70^. For instance, OPCs exposed to demyelination acquire phagocytic and antigen-presenting capabilities, influencing the function of CD8^+^ cytotoxic T cells ^71^ and CD4^+^ helper T cells ^20^ through upregulation of major histocompatibility complex class I (MHC-I)^71^ and MHC-II ^20^ molecules, respectively. However, the prevalence of MHC-expressing oligodendroglial lineage cells is rare ^59^ or absent ^72^ *in vivo*. Moreover, the role of mature OLs, far more abundant than OPCs, in modulating CNS inflammatory and immune responses remains elusive. In this study, we found that SerpinOLs represent a substantial subpopulation of mature oligodendrocytes (40∼60%) during CNS demyelination. They are characterized by gene signatures associated with immune and inflammatory responses and STAT3 signaling activation which underlies SERPINA3N production in OLs. We provide genetic evidence that SerpinOLs perpetuate CNS immune and inflammatory responses to deleterious insults. Thus, our findings point to a potential myelination-independent function of mature OLs through SERPINA3N-regulated neuroinflammation and glial activation.

Our findings challenge the widely cited concept that Serpina3n/SERPINA3N is a reactive astrocyte marker in the diseased CNS. Our results unravel that OLs are the major cell types producing SERPINA3N protein in diverse types of CNS diseases and injuries. A decade ago, *Serpina3n* mRNA was reported as a reliable marker of reactive astrocytes in LPS endotoxic and ischemic brain injury models^35^, and more recently, it has been identified as a representative signature gene of disease-associated astrocytes in Alzheimer’s diseases^37^ and normal aging^37,38^. In contrast, other studies reported that *Serpina3n* is a representative signature gene of disease-associated or specific oligodendrocytes under neurological conditions ^17,18,20,22,33^. Our RNA-seq data reveal that both OLs and astrocytes exhibit comparably high levels of *Serpina3n* mRNA under demyelinating conditions (**Fig. 1l**). However, SERPINA3N protein is produced predominantly in OLs, as confirmed by tdTom reporter mouse line in which tdTom expression depends on SERPINA3N protein translation (**Fig. 2**). These observations led us to hypothesize that *Serpina3n* mRNA is actively translated into SERPINA3N protein in OLs, but the translation is less efficient in astrocytes. Indeed, our RiboTag data (**Fig. 1n**) support this hypothesis, demonstrating that ribosome-bound and actively translated *Serpina3n* mRNA is increased in OLs and much higher than that in astrocytes upon CNS demyelination (**Fig. 1o**). Likewise, in the LPS-induced endotoxic brain model, SERPINA3N protein and tdTom are observed predominantly in OLs (**Fig. 3 c1-c3, Suppl Fig. 6b**) despite high *Serpina3n* mRNA expression in astrocytes ^35^. In Alzheimer’s disease models (5xFAD), our group (**Suppl Fig. 8a, c**) and others ^37^ observed remarkable overlap of SERPINA3N-like signal with GFAP^+^ processes in the subiculum and dentate gyrus, leading to the interpretation that SERPINA3N is a marker of disease-associated astrocytes ^37^. However, we interpreted this observation as imaging artifact rather than true expression because the star-shaped SERPINA3N-like signal does not exist in single IHC using anti-SERPINA3N antibody (**Suppl Fig. 8b, d**). Similarly, in the aged brain, we unequivocally demonstrate that SERPINA3N protein is produced predominantly by OLs using oligodendroglial SERPINA3N cKO mice (*Olig2-Cre:Serpina3n*^fl/fl^). Thus, OLs, but not astrocytes, are the major cell types producing SERPINA3N.

The underlying mechanisms of SerpinOL transition from homeostatic OLs remain poorly defined. SerpinOLs are not specific to CNS demyelinating diseases in which OLs are the direct cellular target. Our data suggest that SerpinOLs may be present in any disease conditions where OL injury occurs. This is supported by our results derived from acute CPZ model. In the CPZ model, approximately 20% of homeostatic OLs already transition to SerpinOLs after a very brief 2-day CPZ treatment when microglial/astroglial activation is yet to be detected. Furthermore, our results collected from genetic mutant mice establish that neuroinflammation which is ubiquitously present in various CNS diseases, are insufficient to drive the state transition of homeostatic OLs into SerpinOLs in the absence of OL injury. Therefore, OL injury, rather than neuroinflammation and glial activation, is a crucial mechanism underlying SerpinOL transition. In this regard, we define SerpinOLs as an injury-transduced activation state and propose their presence as a sensitive marker of OL injury.

While a growing list of disease-associated or specific oligodendroglia is anticipated, their functions are largely unknown ^17^ or speculative ^24^ thus far. The current study provided new insights into the knowledge gaps. By leveraging oligodendroglial SERPINA3N cKO mouse lines ^34^, our results convincingly demonstrate that ablating SERPINA3N attenuates inflammatory responses to CNS demyelination and shifts microglial activation toward a more homeostatic state. To further extend this observation to non-diseased conditions, we used the healthy aging model. Consistently, deleting SERPINA3N remarkably reduces aging-elicited upregulation of proinflammatory cytokines and chemokines and shifts microglial/astroglial activation signatures of aged brains toward those of young brains (**Fig. 9**). Thus, our study provided the first line of evidence suggesting that SerpinOLs, a common and large population of injury-transduced mature OLs, amplify pro-inflammatory responses to CNS diseases and normal aging.

In addition to regulating ubiquitous neuroinflammation, the presence of SerpinOLs in CNS pathologies of distinct etiologies suggests a role of SerpinOLs in regulating disease-specific pathophysiology, such as demyelination, endotoxicity, stroke, CNS trauma, neurodegeneration, and even normal aging. For example, SerpinOLs may actively control extracellular plaque pathophysiology by marking Aβ with SERPINA3N (**Fig. 3f**) in Alzheimer’s disease and animal models. Although SERPINA3N is a secretory protein dysregulated in the body fluids of many neurological conditions ^26^, it is also markedly accumulated and retained in the cytoplasm of injured OLs. The functional consequence of SERPINA3N cytoplasmic accumulation on oligodendroglial survival remains elusive. It has been proposed that SERPINA3N protects oligodendrocytes against injury ^18,73,74^. Our recent *in vitro* study suggests that SERPINA3N may potentiate oligodendrocyte responses to oxidative stress and regulate oligodendrocyte senescence ^34^. Future studies are needed to determine the role of SERPINA3N in oligodendrocyte (or myelin) viability not only in demyelinating diseases but also in other neurological conditions. Our SERPINA3N cKO mouse strains used in the current study will provide powerful and cell-specific tools for this purpose.

We found that SERPINA3N does not influence microglial activation or phenotype polarization in the absence of LPS stimulation *in vitro* (**Fig. 10**). This observation suggests that SERPINA3N is insufficient to trigger glial activation *de novo*. We hypothesize that SERPINA3N exerts its biological functions such as promoting neuroinflammation and glial activation in the context of diseased/injured conditions but not under CNS homeostasis as our recent study concluded ^34^.

This conclusion is in sharp contrast to a recent study ^75^ reporting that SERPINA3N induces robust inflammatory responses in the homeostatic brain. We conjecture that the reported effect in that study ^75^ may result from CNS trauma elicited by injection needles during experimenting. In this regard, homeostatic transgenic mice carrying enforced *Serpina3n*-expressing alleles will be required to prove or falsify this hypothesis. Furthermore, it would be interesting to define the role of SerpinOLs in CNS traumatic injury (**Suppl Fig. 9**) using our cell-specific genetic mouse models.

The molecular mechanisms underlying SERPINA3N-regulated microglial activation remain to be established. SERPINA3N inhibits the cleavage activity of endogenous serine proteases. Yet, other functions beyond its protease inhibition are not uncommon ^26^. At the molecular level, SERPINA3N is secretory glycoprotein with N-linked glycosylation which could be recognized by microglia through “yet-to-be-identified” microglial receptors to modulate their activation and function ^76^. Therefore, identifying SERPINA3N binding partners (serine proteases and non-protease receptors) will provide novel molecular insights into how SERPINA3N regulates microglial activation and function.

To summarize, our findings nominate SerpinOLs as a subpopulation of injury-transduced OLs. Direct injury to OLs, rather than neuroinflammation, underlies the state transition of SerpinOLs. Phenotypically, SerpinOLs acquires molecular signatures of inflammatory and immune regulation. We also found that STAT3-mediated signaling is activated in SerpinOLs. Functionally, SerpinOLs perpetuate neuroinflammation and promote microglia toward pro-inflammatory and disease-associated states. Thus, targeting SerpinOLs and their transcriptomic profiles represents a promising avenue modulating CNS inflammatory response to various CNS pathologies. Future studies are needed to define if and how STAT3 activates Serpina3n expression and the cell-intrinsic role of SERPINA3N dysregulation in oligodendrocyte survival or damage.

## 12 Supplementary Figures and Legends

**Supplementary Figure 1.**
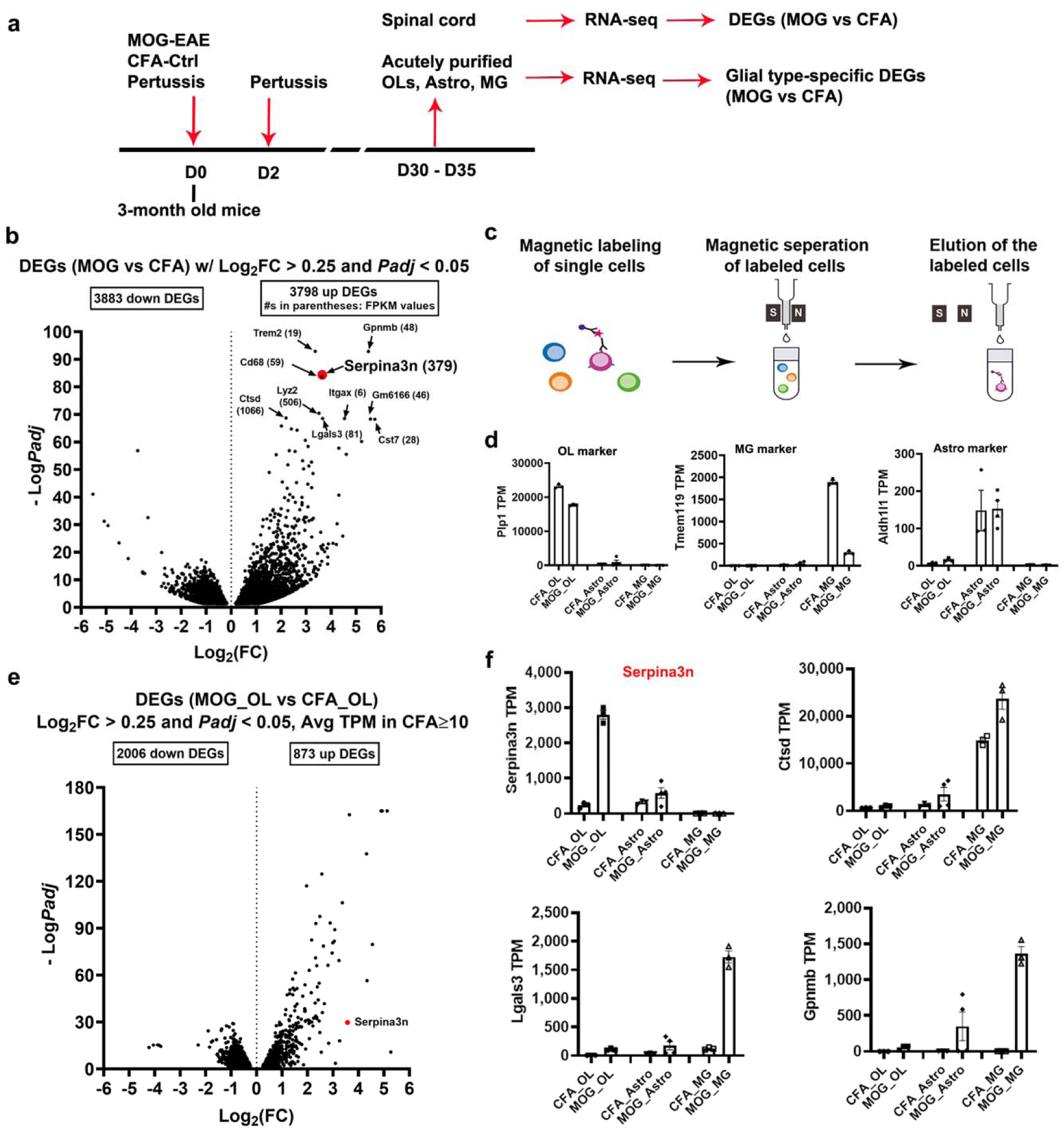
Identification of glial population-specific differentially expressed genes (related to Figure 1). **a** Experimental design. Inflammatory demyelination of experimental autoimmune encephalomyelitis (EAE) was induced in adult mice via MOG immunization (CFA as control) at day 0 (D0), with pertussis administered i.p. on D0 and D2. RNA-seq was performed on whole spinal cords (**b**) and MACS-purified glial populations (**c-f**) at D30-D35. OLs, oligodendrocytes; Astro, astrocytes; MG, microglia. **b** Volcano plot of differentially expressed genes (DEGs) in spinal cord of MOG/EAE (n=4 mice) vs. CFA mice (n=3 mice) at D35. Top10 DEGs are labeled with gene symbols and FPKM values. **c** Schematic of magnetic beads-assisted cell sorting (MACS) of spinal cord glia at D30. OL, Astro, and myeloid cells (MG/macrohages) were isolated using anti-O4, anti-ACSA2, and anti-CD11b magnetic beads, respectively. **d** Purity validation of sorted population using representative markers *Plp1* (OL), *Tmem119* (MG), and *Aldh1l1* (Astro). Expression is shown as TPM (transcripts per million). *n* = 3 mice per group for CFA_OL, MOG_OL, CFA_Astro, CFA_MG and MOG_MG; *n* = 4 mice for MOG_Astro. **e** Volcano plot of DEGs in MOG_OL versus CFA_OL. *Serpina3n* is highlighted in red. **f** TPM expression of *Serpina3n*, *Ctsd*, *Lgals3*, and *Gpnmb* in OLs, Astro, and MG from MOG/EAE and CFA mice. *Serpina3n* is specifically upregulated in OL, while *Ctsd*, *Lgals3*, and *Gpnmb* are enriched and upregulated in MG. *n* = 3 mice per group for CFA_OL, MOG_OL, CFA_Astro, CFA_MG and MOG_MG; *n* = 4 mice MOG_Astro. Data were presented as mean ± s.e.m. (d, f).

**Supplementary Figure 2.**
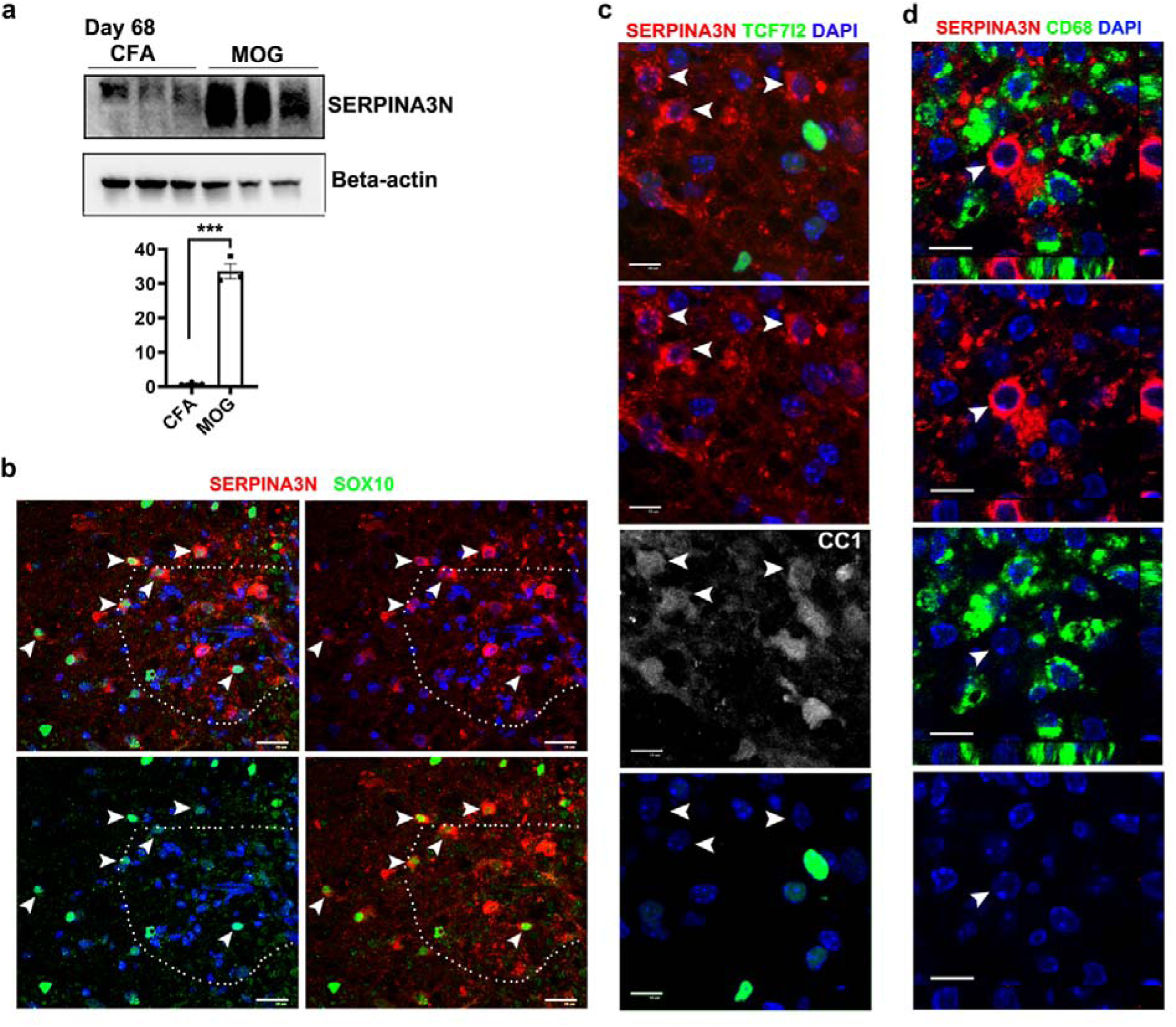
SERPINA3N persists into chronic phase of inflammatory demyelination (related to Figure 1). **a** Western blot showing SERPINA3N levels in spinal cords. *** *P* = 0.0001. *n* = 3 mice per group (error bars, s.e.m.). **b** Immunofluorescence of SERPINA3N and SOX10 in MOG/EAE spinal cord. Dotted regions indicate inflammatory lesions with dense DAPI^+^ nuclei. Arrowhead, SERPINA3N^+^SOX10^+^ cells. **c** Confocal images of SERPINA3N, CC1, and newly regenerated OL marker TCF7l2 in MOG/EAE spinal cord. Evenly distributed DAPI^+^ cells indicate non-lesional areas. Arrowheads highlight SerpinOLs lacking TCF7l2 expression. **d** Orthogonal view of confocal images of SERPINA3N and the activate myeloid cell marker CD68. Arrowhead points to a SerpinOL adjacent to CD68^+^ cells. All panels: D68 post-MOG (or CFA). Nuclei counterstained with DAPI (blue), Scale bars: **b** 20 µm, **c, d** 10 µm.

**Supplementary Figure 3.**
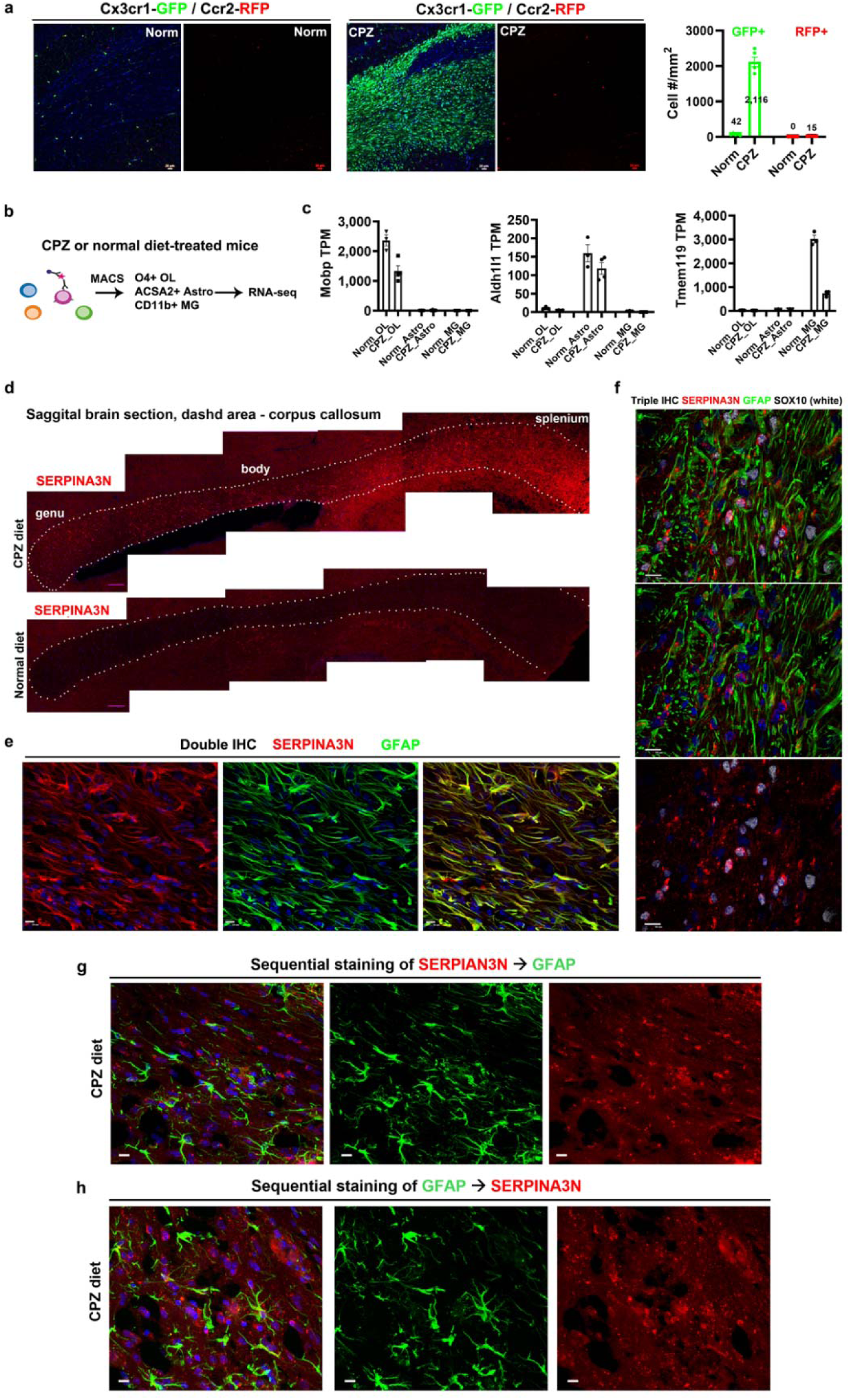
SERPINA3N expression in the corpus callosum during cuprizone (CPZ)-induced demyelination (related to Figure 1). **a** Fluorescent IHC and quantification of GFP^+^ (resident microglia and/or peripheral monocyte-derived macrophages) and RFP^+^ (peripheral monocyte-derived macrophages) in the corpus callosum of Cx3cr1-GFP/Ccr2-RFP mice after 4 weeks CPZ or Norm diet. CPZ-induced demyelination showed robust GFP^+^/RFP^-^ microglial expansion and rare RFP^+^ infiltrated myeloid cells. DAPI counterstaining (blue). Scale bars: 20 µm. *n* = 5 mice per group (error bars, s.e.m.). **b-c** MACS purification of brain OLs, Astro, and MG of normal (Norm) or 4 weeks CPZ diet (**b**) and purity assessment using lineage-specific markers (*Mobp* for OLs, *Aldh1l1* for Astro, *Tmem119* for MG) (**c**). TPM (transcripts per million) values are from RNA-seq. *n* = 3 mice per group for Norm_OL, Norm_Astro, Norm_MG and CPZ_MG; *n* = 4 mice per group for CPZ_OL and CPZ_Astro (error bars, s.e.m.). **d** Single fluorescent IHC showing SERPINA3N induction along the splenium-to-genu gradient in CPZ (4 wks) corpus callosum (dotted regions), with no or little expression in Norm mice. Scale bars: 200 µm. **e** Double fluorescent IHC showing near-complete overlap of SERPINA3N with reactive astrocyte marker GFAP in 4 weeks CPZ corpus callosum. DAPI counterstaining (blue). Scale bars: 10 µm. **f** Triple fluorescent IHC showing no overlap of SERPINA3N with GFAP, but co-localization with oligodendroglial marker SOX10 in CPZ (4 wks) corpus callosum. DAPI counterstaining (blue). Scale bars: 10µm. Note the distinct SERPINA3N staining patterns in panel **e** vs. **f**. **g-h** Sequential IHC in CPZ diet corpus callosum with reversed staining order to validate SERPINA3N-GFAP signal overlap. SERPINA3N staining performed first, followed by GFAP (**g**). GFAP staining performed first, followed by SERPINA3N (**h**). Scale bars: 10 µm.

**Supplementary Figure 4.**
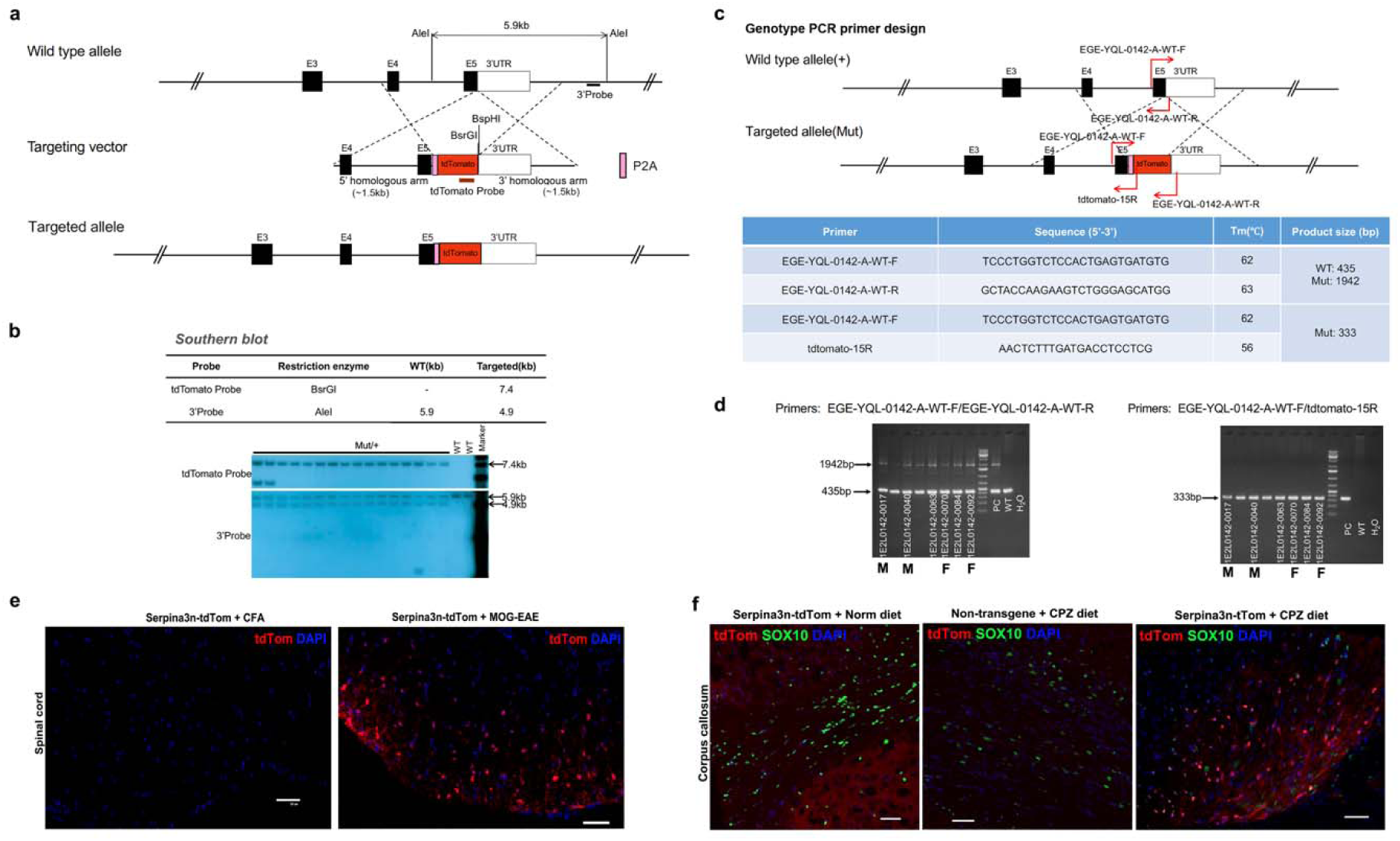
Generation of Serpina3n-tdTomato knock-in reporter mice (related to Figure 2). **a** Schematic of the wild-type (WT) *Serpina3n* allele and targeting construct. The *Serpina3n* stop codon was replaced with a *P2A-tdTom* sequence, allowing expression of separate SERPINA3N and tdTom proteins. Southern Blot probe locations (tdTom probe and 3’ probe) are indicated. **b** Southern Blot confirming successful *Serpina3n-tdTom* integration F1 mice. **c** Genotyping primer sequences and expected PCR product sizes. The EGE-YQL-0142-A-WT-F/tdtomato-15R pair amplifies a 333 bp product from the mutant allele; EGE-YQL-0142-A-WT-F/EGE-YQL-0142-A-WT-R amplifies a 435 bp WT product and a 1942 bp mutant product. **d** Agarose gel showing PCR products for T and mutant alleles. **e** Fluorescent IHC of tdTom in the spinal cord of Serpina3n-tdTom mice at D30 post-MOG/EAE (or CFA). Note that tdTom is induced by MOG/EAE but absent in CFA controls. Scale bar: 50 µm. **f** Fluorescent IHC of tdTom in the corpus callosum of Serpina3n-tdTom mice or non-transgenic mice after 4-week CPZ or normal (Norm) diet. tdTom is detected only in CPZ-treated Serpina3n-tdTom mice. Scale bar: 50 µm.

**Supplementary Figure 5.**
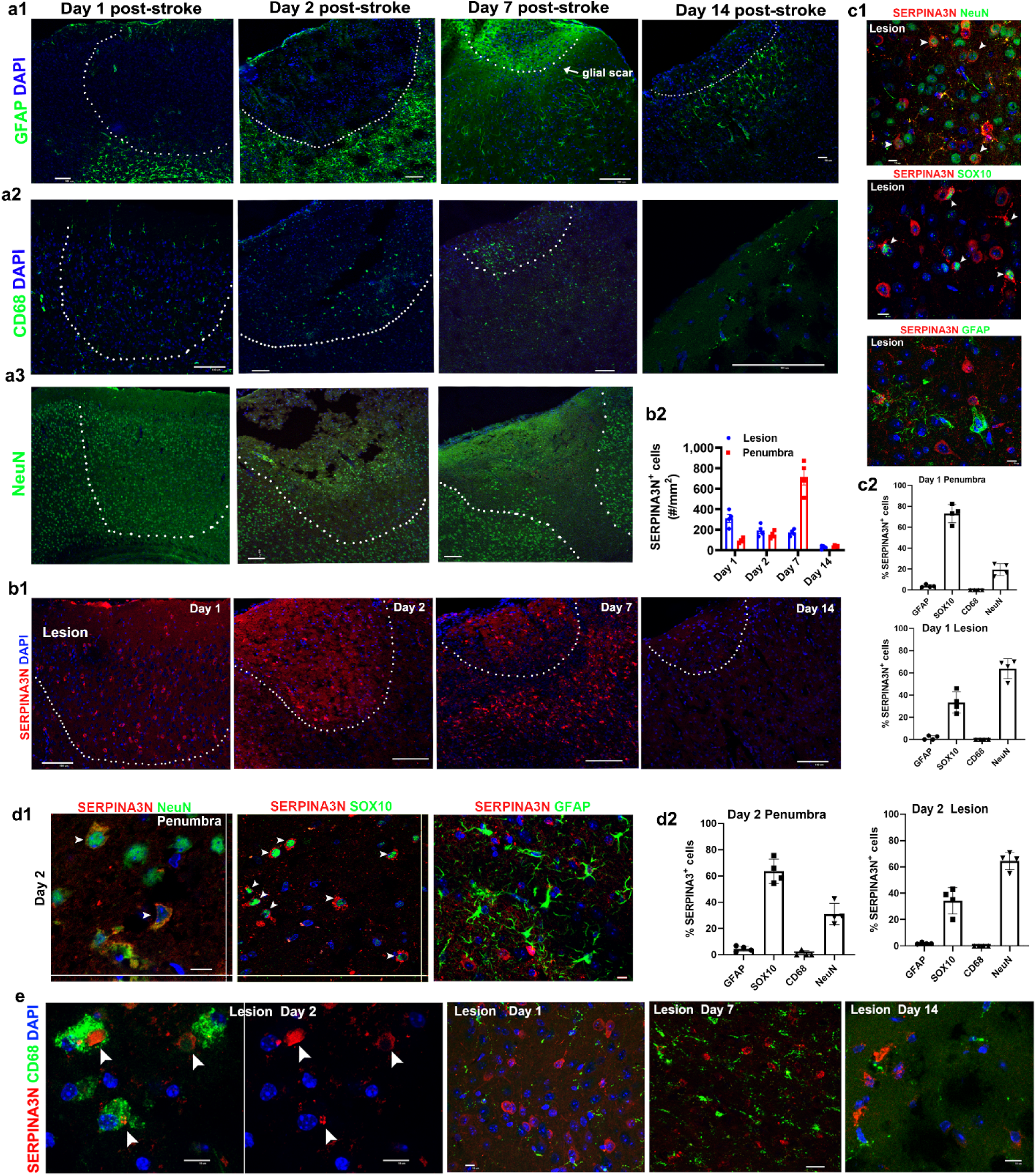
SERPINA3N expression and cellular identity in photothrombotic stroke (related to Figure 3). **a1-a3**, temporal expression of GFAP (reactive astrocytes), CD68 (microglial activation), and NeuN (neuronal loss) following photothrombotic stroke. Scale bars: 100 µm. **b1-b2**, densities of SERPINA3N^+^ cells in lesion cores (dotted) and penumbra at at day 1, 2, 3, and 14 post-stroke injury. n = 4 mice per group (mean ± s.e.m.). Scale bars: 100 µm. **c1-c2**, representative IHC images and quantification of SERPINA3N^+^ cells at D1 post-stroke injury. n = 4 mice per group (mean ± s.d.). Scale bars: 10 µm. **d1-d2**, representative IHC images and quantification of SERPINA3N^+^ cells at D2 post-stroke injury. n = 4 mice per group (mean ± s.d.). Scale bars: 10 µm. **e**, fluorescent IHC showing CD68^+^ myeloid cells engulfing extracellular SERPINA3N (arrowheads) but lacking SERPINA3N expression during the acute phase. Scale bars: 10 µm.

**Supplementary Figure 6.**
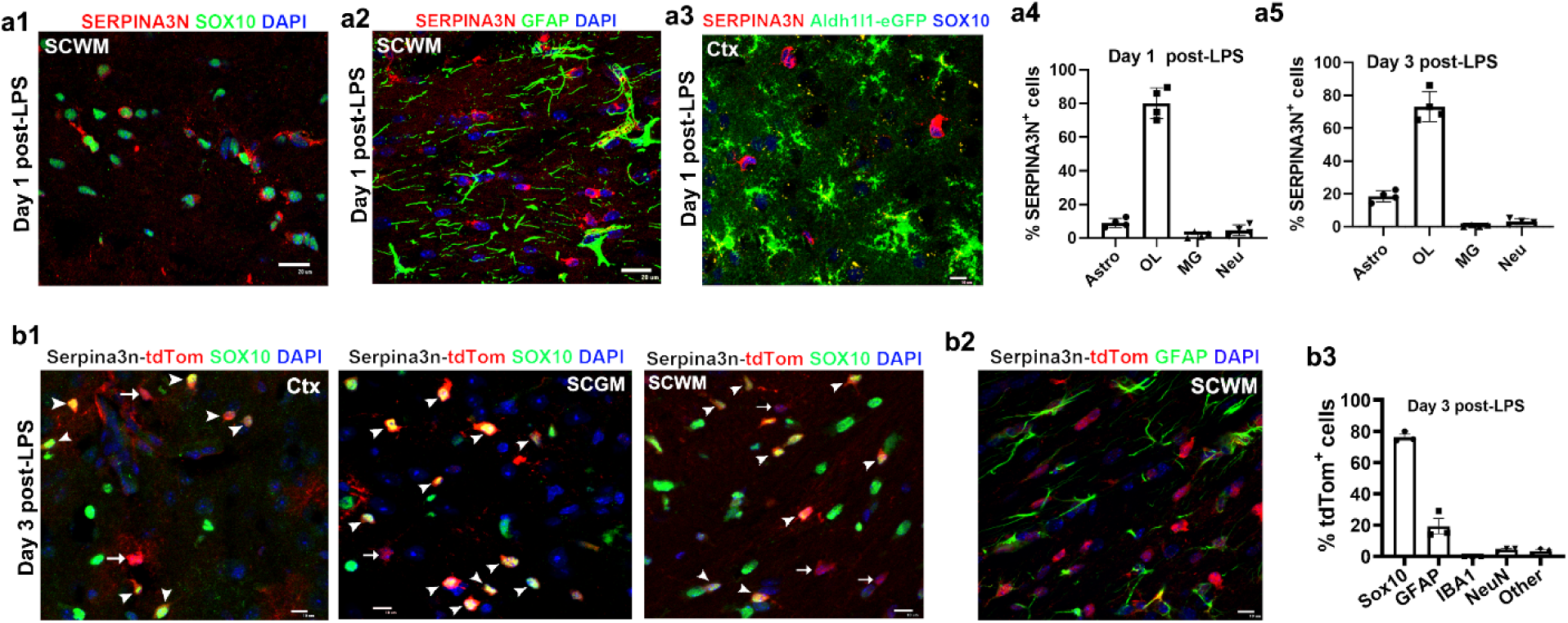
SERPINA3N expression and cellular identity in LPS-induced endotoxicity (related to Figure 3). **a1-a2**, double fluorescent IHC of SERPINA3N with SOX10 or GFAP in subcortical white matter (SCWM) at D1 post-LPS injection. Scale bars: 10 µm. **a3**, triple fluorescent IHC of SERPINA3N, SOX10 and Aldh1l1-GFP (astrocyte reporter) in cortex (Ctx) at D1 post-LPS injection. Scale bar: 10 µm. **a4-a5**, percentages of SERPINA3N^+^ cells expressing GFAP (Astro), SOX10 (OLs), IBA1 (MG), or NeuN (Neurons) at D1 and D3 post-LPS injection. n = 4 mice each group (error bars, s.d.). Scale bar: 20 µm. **b1-b3**, double fluorescent IHC of Sepina3n-tdTom reporter with SOX10 or GFAP in Ctx, subcortical gray matter (SCGM), and subcortical white matter (SCWW), and quantification at day 3 post-LPS. n = 4 mice each group (error bars, s.d.). Scale bar: 10 µm.

**Supplementary Figure 7.**
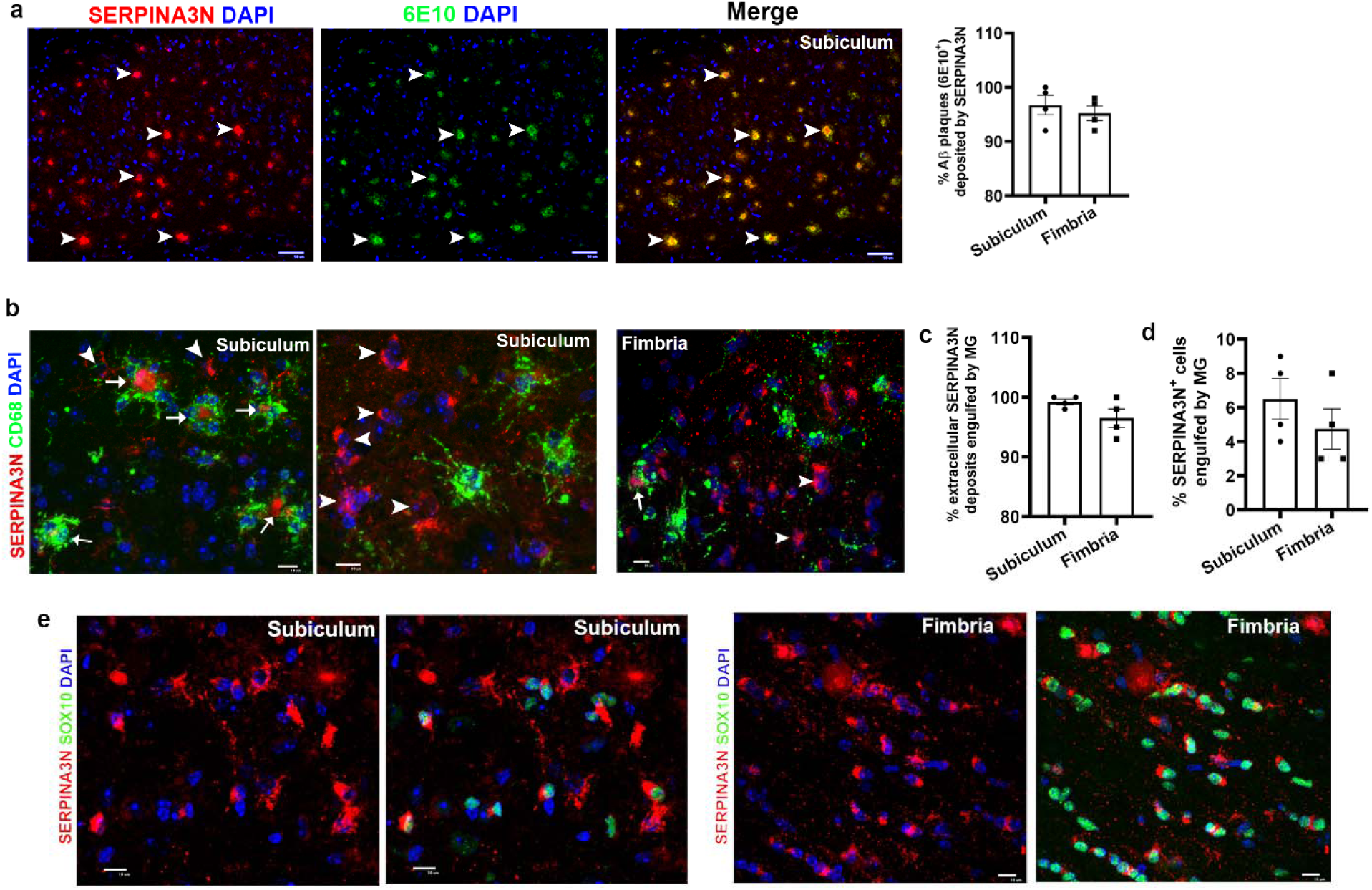
association of extracellular and intracellular SERPINA3N with amyloid beta plaques, microglia, and oligodendroglia (related to Figure 3). **a**, double IHC showing co-localization of extracellular SERPINA3N (arrowheads) with β-amyloid plaques (6E10). n = 4 mice (error bars, s.e.m.). Scale bar: 50µm. **b**, double IHC showing engulfment of extracellular SERPINA3N (arrows) by CD68^+^ microglia; intracellular SERPINA3N (arrowheads) was not engulfed. Scale bar: 10µm. **c-d**, quantification of extracellular (**c**) and intracellular (**d**) SERPINA3N engulfed by microglia. n = 4 mice (error bars, s.e.m.). **e**, double IHC showing intracellular SERPINA3N co-localized with SOX10^+^ oligodendrocytes. Scale bars: 10 µm. 7-8-month-old 5xFAD transgenic mice were used.

**Supplementary Figure 8.**
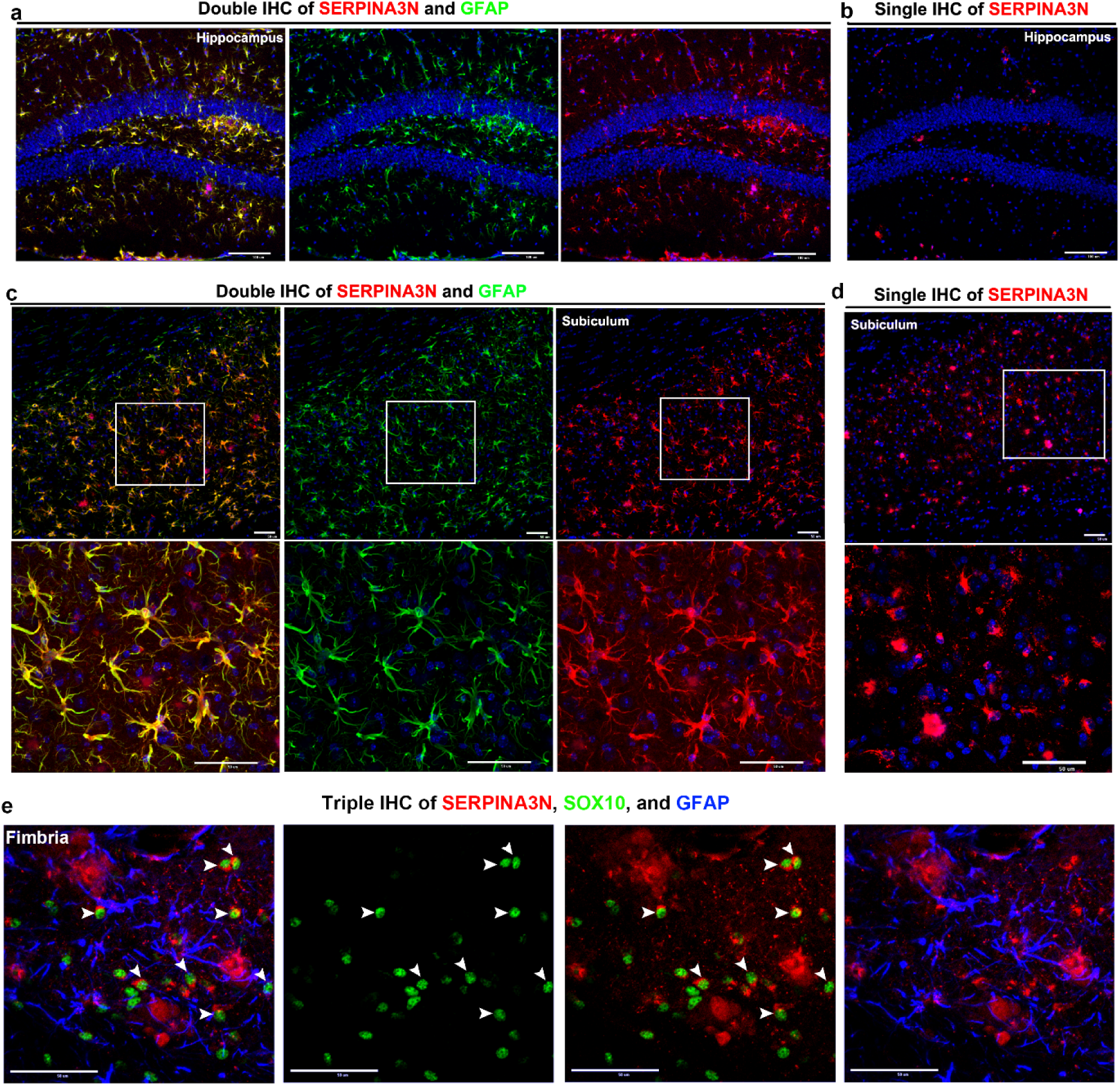
evidence of SERPINA3N immunoreactivity artifacts in GFAP^+^ astrocytes (related to Figure 3). **a-b,** double IHC for SERPINA3N and GFAP (**a**) and single IHC for SERPINA3N (**b**) on adjacent hippocampal sections of 5xFAD mice. SERPINA3N signal overlaps with GFAP^+^ astrocytes in double IHC but shows distinct patterns in single IHC. Blue, DAPI^+^ nuclei. **c-d,** double IHC for SERPINA3N and GFAP (**c**) or single IHC for SERPINA3N (**d**) in adjacent sections of the subiculum of 5xFAD mice. Boxed areas shown at higher magnification. As in panels A-B, SERPINA3N signal overlaps with GFAP in double IHC but not in single IHC. Blue, DAPI⁺ nuclei. **e,** triple IHC of SERPINA3N, GFAP, and SOX10 in the hippocampal fimbria. SERPINA3N co-localizes with SOX10^+^ cells (arrowheads) but not GFAP. 8-month-old 5xFAD mice were used. Scale bars: 100 µm (**A–B**); 50 µm (**C–E**).

**Supplementary Figure 9.**
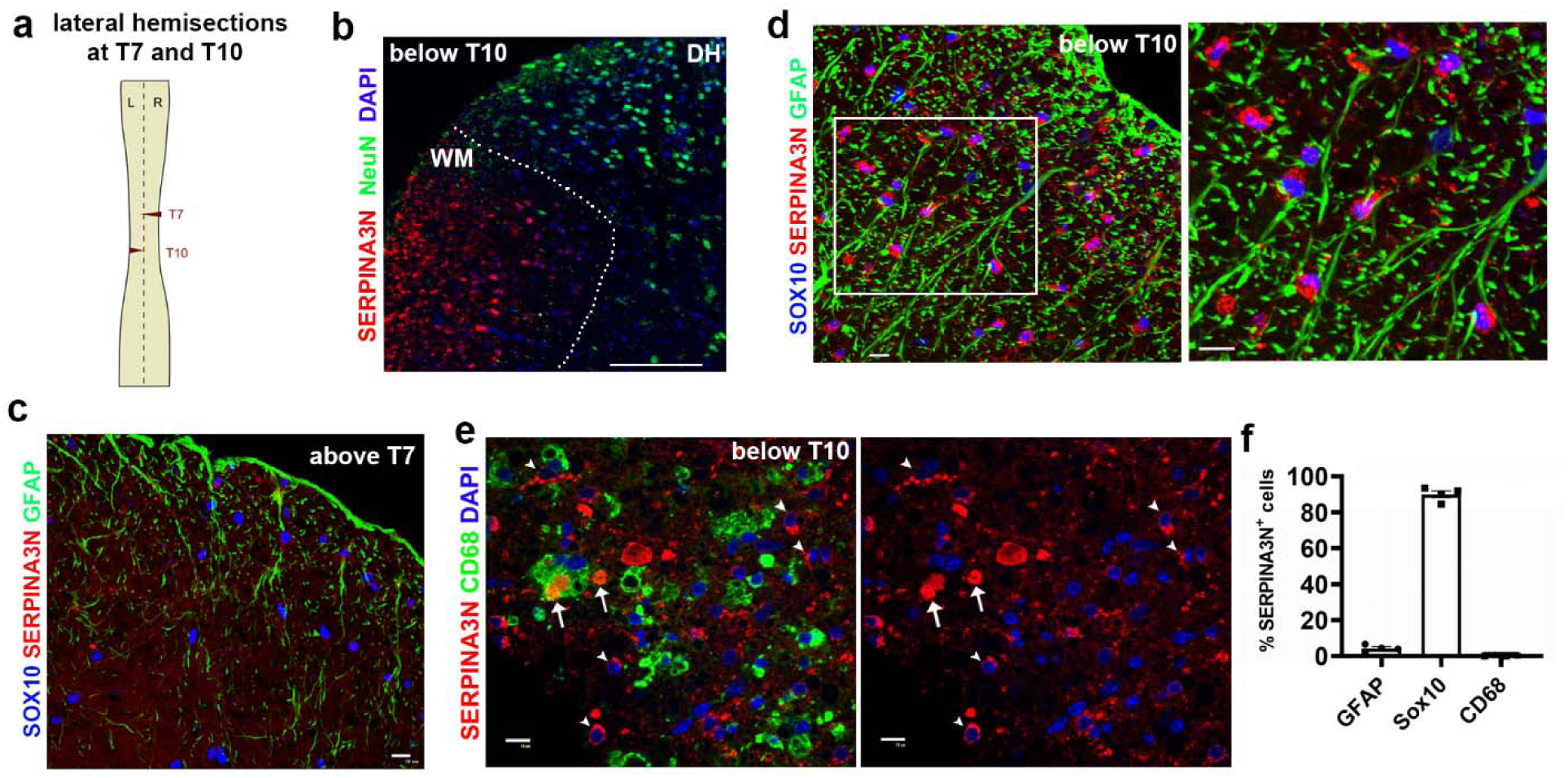
oligodendrocytes transition into SerpinOLs in response to CNS trauma (related to Figure 3). **a**, lateral spinal cord hemisection at thoracic (T) segments T7 and T10, fully disrupting innervation below T10. **b**, SERPINA3N is induced in white matter (WM) but not in gray matter regions containing NeuN⁺ neurons. DH, dorsal horn. **c-d**, triple IHC for SERPINA3N, SOX10, and GFAP in spinal cord segments above T7 (**c**) and below T10 (**d**). Boxed area in **d** shown at higher magnification (right). **e**, extracellular SERPINA3N (arrows) is engulfed by CD68⁺ myeloid cells; intracellular SERPINA3N (arrowheads) is not. **f**, percentage of SERPINA3N⁺ cells co-expressing indicated markers below T10. n = 3 mice (error bars, s.e.m.). Mice were analyzed 10 weeks post-SCI. Scale bars: 10 µm.

**Supplementary Figure 10.**
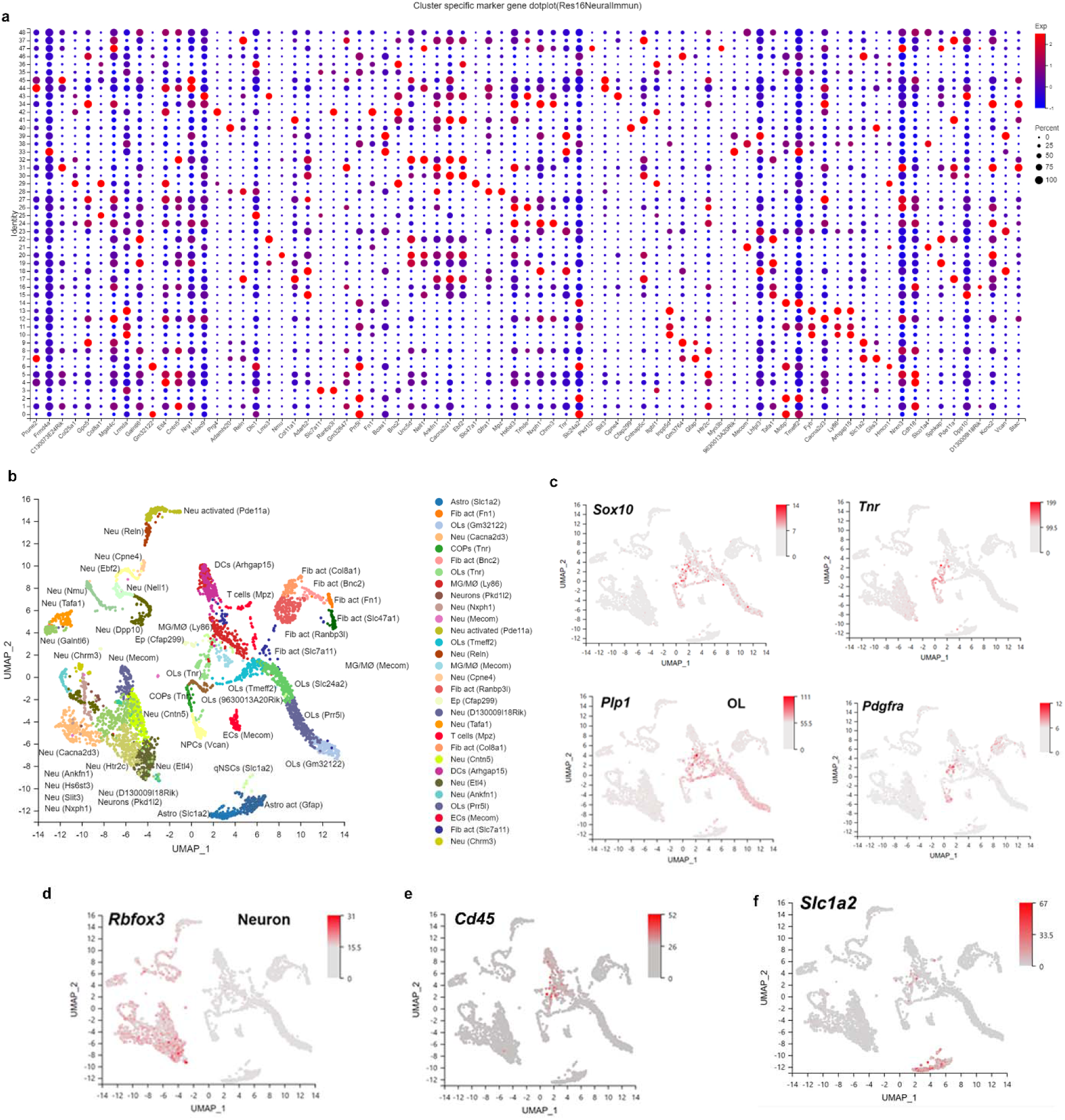
lineage-specific marker expression in UMAP clusters of scRNA-seq (related to Figure 7). **a,** dot plot visualizing the expression of canonical marker genes used to assign cell identity to each scRNA-seq cluster. The size of the dot represents the percentage of cells expressing the gene, and the color intensity represents the average expression level (red = high, blue = low). **b,** cell type annotations and representative marker genes. scRNA-seq were performed on D30 MOG spinal cord. **c**, expression levels and distribution of stage-dependent oligodendroglial lineage marker genes, *Sox10, Tnr, Plp1,* and *Pdgfra.* **d**, expression levels and distribution of the neuronal marker gene *Rbfox3* (NeuN). **e**, expression levels and distribution of the immune cell marker gene *Cd45*. **f**, expression levels and distribution of the astrocyte marker gene *Scl1a2* (glutamate transporter 1, GLT1).

**Supplementary Figure 11.**
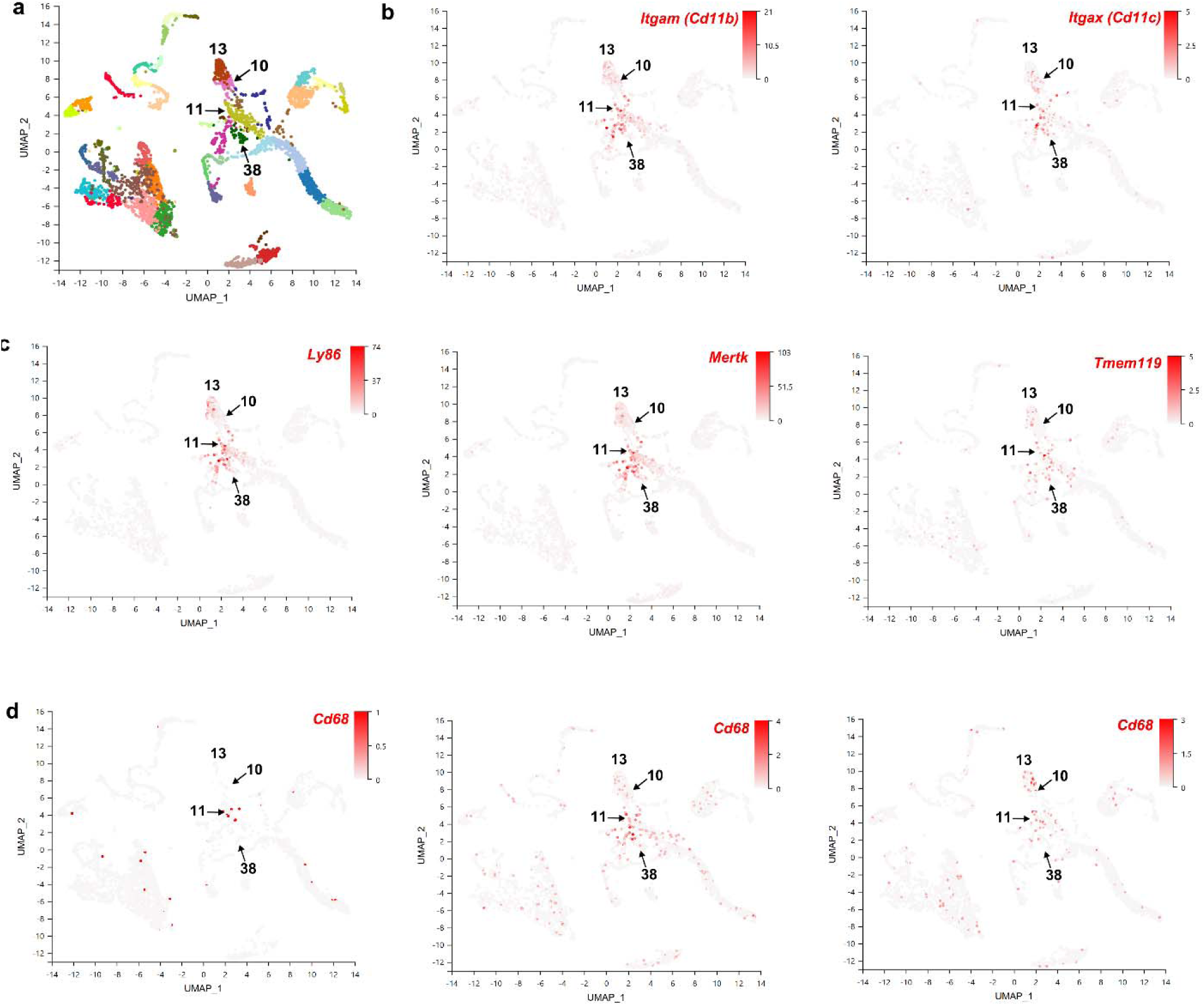
myeloid cell clusters of scRNA-seq of MOG/EAE (related to Figure 8). **a**, cluster distribution of myeloid cells. Cluster 11, 13, and 38 were annotated as microglia/macrophages (MG/MØ), and cluster 10 as dendritic cells (DCs). **b**, distribution and expression level of myeloid cell gene *Cd11b* and DC-enriched gene *Cd11c*. **c**, distribution and expression level of activated myeloid cell genes *Ly86* and *Mertk*, and microglial gene *Tmem119*. **d**, distribution and expression level of the phagocytotic cell marker *Cd68* in CFA (left), Ctrl_MOG (middle), and Serpina3n cKO_MOG (right) mice.

**Supplementary Figure 12.**
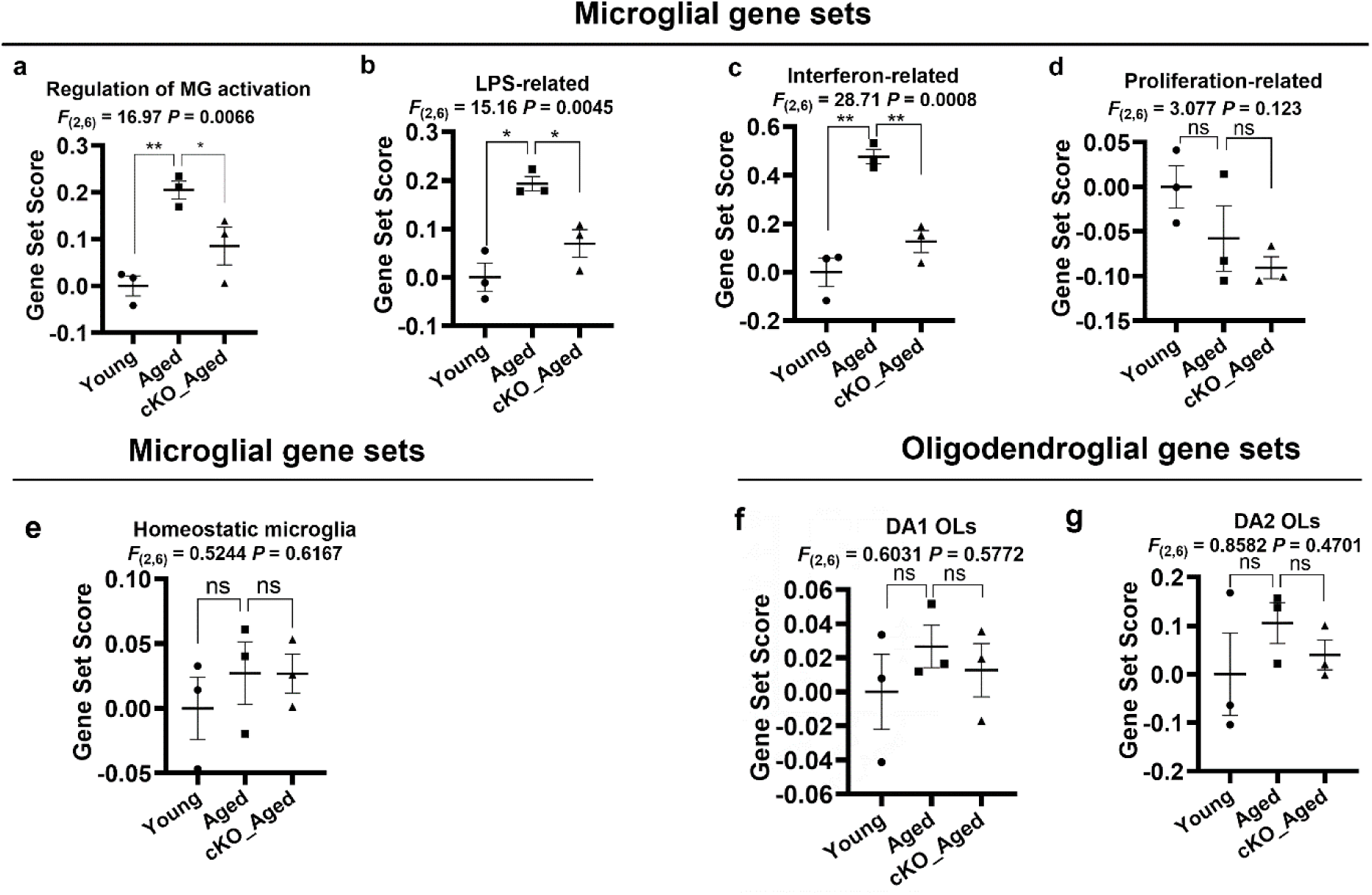
Ablating SERPINA3N alters the activation state of microglia and oligodendrocytes at the transcriptomic level (related to Figure 9). **a**, cumulative gene set scores of microglial activation regulation (Suppl Table 11) in each group of mice. One way ANOVA, followed by Holm-Sidak multiple comparisons test. *P* = 0.0069 Aged vs Young, *P* = 0.0497 cKO_Aged vs Aged. **b-d**, cumulative scores of microglial gene sets ^62^ involved in LPS-stimulation (**b**), interferon-response (**c**), and proliferation (**d**) (Suppl Table 11). One way ANOVA followed by Holm-Sidak multiple comparisons test. LPS *P* = 0.0048 Aged vs Young, *P* = 0.0236 cKO_Aged vs Aged; Interferon *P* = 0.0.0010 Aged vs Young, *P* = 0.0034 cKO_Aged vs Aged; Proliferation *P* = 0.3090 Aged vs Young, *P* = 0.4102 cKO_Aged vs Aged. **e**, gene set scores of microglial homeostasis markers ^77^ (Suppl Table 11) of young (2 mon), aged (20 mon), and Serpina3n cKO aged (20 mon) mice. One way ANOVA followed by Holm-Sidak multiple comparisons test. *P* = 0.7911 Aged vs Young, *P* = 0.9918 cKO_Aged vs Aged. **f-g**, cummulative gene set scores of DA1 OLs (associated with immunogenic genes) and DA2 OLs (associated with cell survival and differentiation genes) (Suppl Table 11). One way ANOVA followed by Holm-Sidak multiple comparisons test. DA1 OLs *P* = 0.6777 Aged vs Young, *P* = 0.8288 cKO_Aged vs Aged; DA2 OLs *P* = 0.5649 Aged vs Young, *P* = 0.6974 cKO_Aged vs Aged. n = 3 mice each group (error bars stand for s.e.m.)

## 12 Supplementary Tables and 6 GEO Datasets

**Suppl Table 1** - Expression levels (FPKM) of differentially expressed genes (DEGs) in the spinal cord at D36 post-EAE (related to Fig. 1).

**Suppl Table 2** - Glial type-enriched differentially expressed genes (DEGs) in response to MOG-EAE injury (related to Fig. 1).

**Suppl Table 3** - RNA-seq of MACS-purified MG, OL and Astro in CPZ-treated brain (related to Fig. 1)

**Suppl Table 4** - DEGs and functional annotations in the spinal cord of CFA versus PBS (related to Fig. 5).

**Suppl Table 5** - DEGs and functional annotations in the optic nerve of NFkB CA versus NFkB Ctrl (related to Fig. 5).

**Suppl Table 6** - Marker genes used for UMAP clustering of EAE spinal cord scRNA-seq (related to Fig. 7).

**Suppl Table 7** - DEGs and functional annotation of SerpinOLs versus non-SerpinOLs in MOG-EAE (related to Fig. 7).

**Suppl Table 8** - Pseudobulk DEGs and functional annotations of microglia and macrophages in the spinal cord of Ctrl_MOG vs Ctrl_CFA (related to Fig. 8).

**Suppl Table 9 -** Pseudobulk DEGs and functional annotations of microglia and macrophages in the spinal cord of cKO_MOG vs Ctr_MOG (related to Fig. 8).

**Suppl Table 10 -** DEGs in microglia and GO terms of CPZ_cKO, CPZ_Ctrl, and Norm_Ctrl mice (4 weeks CPZ) (related to Fig. 8).

**Suppl Table 11 -** List of gene sets and gene symbols in each gene set (related to Fig. 9 and Fig. 9).

**Suppl Table 12 –** DEGs in the brain of young, aged, and cKO-aged mice (related to Fig. 9).

**GSE287324** (Suppl Table 1 and Suppl Table 4), **GSE287323** (Suppl Table 2); **GSE287320** (Suppl Table 3); **GSE287310** (Suppl Table 6, Suppl Table 7, Suppl Table 8, and Suppl Table 9); **GSE287313** (Suppl Table 10); **GSE287309** (Suppl Table 12).

## Materials and Methods

### Transgenic mice

All transgenic mice were maintained on a C57BL/6 background and housed in 12 h light/dark cycle with water and food, and both males and females were used in this study. Transgenic lines used in the study, including *Serpina3n-tdTom* reporter line we generated by our own group, *Plp-CreER^T^*^2^ (RRID:IMSR_JAX:005975), *Aldh1l1-CreER^T^*^2^ (RRID:IMSR_JAX:031008), *Aldh1l1-eGFP* (RRID:IMSR_JAX:030247), *RiboTag* (RRID:IMSR_JAX:029977), *Olig2-Cre* (RRID:IMSR_JAX:025567), *Serpina3n-floxed* (RRID:IMSR_JAX:027511), *5xFAD* (RRID:MMRRC_034848-JAX), *Cx3cr1-GFP/Ccr2-RFP* (RRID:IMSR_JAX:032127), and *Cnp-Cre:R26Stop^FL^ikk2ca* (referred to as NFkB CA mice) was from our previous study ^46^.

Serpina3n-tdTom knock-in reporter mouse line was generated using CRISPR/Cas9-based Extreme Genome Editing (EGE) technology. A single sgRNA was designed to target exon 5 of the mouse *Serpina3n* gene. The targeting construct was assembled using the TV-4G vector and included a ∼1.5 kb 5’ homologous arm, exons 4–5 fused with a P2A-tdTomato cassette, and a ∼1.5 kb 3’ homologous arm. The targeting vector DNA, Cas9 mRNA, and sgRNA were microinjected into zygotes derived from C57BL/6J mice. The injected embryos were subsequently transferred into pseudopregnant female mice. The P2A-tdTomato cassette was precisely inserted into exon 5, immediately upstream of the 3’ untranslated region (3’UTR) of the *Serpina3n* gene. F0 positive founder mice were bred with wild-type C57BL/6J mice to obtain F1 offspring. Germline transmission in F1 heterozygous mice was confirmed by PCR, DNA sequencing, and Southern blot analysis using probes at both the 5’ and 3’ ends to exclude random integration events. The DNA template for PCR was prepared using: EGE-YQL-0142-A-WT-F: TCCCTGGTCTCCACTGAGTGATGTG; EGE-YQL-0142-A-WT-R: GCTACCAAGAAGTCTGGGAGCATGG and tdtomato-15R: AACTCTTTGATGACCTCCTCG (see Suppl Figure 4).

Animal genotype was determined by PCR of genomic DNA extracted from tail tissue. All Cre lines were maintained as heterozygosity. All animals and procedures were approved by UC Davis IACUC.

### Mouse models of CNS diseases and injuries

#### MOG/EAE model

Both male and female mice were used in this study. 12–15-week-old mice were EAE-induced using two subcutaneous flank injections of a total of 300 µg of MOG peptide (35-55) emulsified in complete Freund’s adjuvant [incomplete Freund’s adjuvant (ThermoFisher) containing 5 mg/ml of heat-killed Mycobacterium tuberculosis (BD Difco Adjuvants)]. On days 0 and 2 after induction, mice also received intraperitoneal injections of 200 ng of pertussis toxin. CFA control mice received only CFA and pertussis toxin, without MOG peptide. Mice were weighed and assessed for clinical symptoms for classical EAE on a 5-point scale using our published protocols^79^.

#### Cuprizone (CPZ) model

Mice were fed 0.25% (w/w) CPZ (bis-cyclohexanone oxaldihydrazone, Inotiv, #TD.140805) mixed into standard powdered rodent chow for 2 days to 4 weeks to induce demyelination. Food consumption was monitored and changed twice weekly. Control mice were fed standard powdered mouse chow for up to 4 weeks. Mice were euthanized after 2 days and 4 weeks of CPZ treatment, and brain tissues were collected for cell sorting, molecular analyses, and various histological examinations.

#### LPS endotoxicity

As previously describe^35^, 6-8 weeks old mice were injected intraperitoneally (i.p.) with 3 mg/kg of LPS (Cat# L2880, *E. coli* O55:B5, Sigma Aldrich, USA) or PBS (Cat# 10010023, Gibco, USA) for endotoxin challenge.

#### Photothrombosis-induced focal cerebral ischemia

We used our published protocol to induce photothrombotic focal cerebral ischemia^39^. In brief, Rose Bengal (RB), a photosensitive dye, was dissolved in artificial cerebrospinal fluid (ACSF) and administered via tail vein injection at a concentration of 0.03 mg/g body weight. For photoactivation, a defined region at the center of the craniotomy was exposed to green light (535 ± 25 nm) emitted from a mercury lamp for 2 minutes using a 10× objective lens with a numerical aperture of 0.3.

#### 5xFAD mouse model for Alzheimer’s disease

To investigate SERPINA3N expression in the context of Alzheimer’s disease, we utilized 7-8 months old hemizygous 5XFAD transgenic mice, which are a widely used model of aggressive amyloid pathology. These mice overexpress mutant human amyloid beta (A4) precursor protein 695 (APP) harboring three Familial Alzheimer’s Disease (FAD) mutations, Swedish (K670N/M671L), Florida (I716V), and London (V717I), along with human presenilin 1 (PSEN1) transgene carrying two additional FAD mutations (M146L and L286V). 5XFAD mice were obtained from the Mutant Mouse Resource and Research Center (MMRRC) (RRID: MMRRC_034848-JAX) and maintained on a C57BL/6J background by crossing hemizygous 5XFAD mice with wild-type C57BL/6J mice (RRID: IMSR_JAX:000664). Genotyping of the 5XFAD transgene, located on mouse chromosome 3, was performed using the following primers: common forward 5′-ACC CCC ATG TCA GAG TTC CT-3′, wild-type reverse: 5′-TAT ACA ACC TTG GGG GAT GG-3′, and mutant reverse: 5′-CGG GCC TCT TCG CTA TTA C-3′. PCR amplification yields a 129 bp band for the transgene and a 216 bp band for the wild-type allele. Heterozygous mice exhibit both bands, whereas wild-type mice show only the 216 bp band.

#### Spinal cord injury

Spinal cord injury of T7 and T10 double lateral hemisection was described in our previous study^80^. Briefly, a midline incision was made over the thoracic vertebrae, followed by a T7–10 laminectomy. For the T7 right side over-hemisection, we carefully used both a scalpel and micro-scissors to interrupt the bilateral dorsal column at T7 and ensured no sparing of ventral pathways on the contralateral side (Figure S9A). For the T10 left hemisection, we carefully used both a scalpel and micro-scissors to interrupt only the left side of the spinal cord until the midline. The muscle layers were then sutured, and the skin was secured with wound clips. All animals were sacrificed 10 weeks after surgery.

#### Normal aging

To determine whether OLs are a source of SERPINA3N during normal aging, we generated transgenic *Olig2-Cre:Serpina3n*^fl/fl^ mice (oligodendroglial *Serpina3n* cKO). We analyzed three experimental groups: young control mice (2 months old), aged wild-type mice (20 months old), and aged Serpina3n cKO mice (20 months old, *Olig2-Cre:Serpina3n*^fl/fl^). Mice were euthanized at the designated ages, and their forebrains were dissected and collected for downstream histological, molecular, and biochemical analyses.

#### Tissue preparation, immunohistochemistry (IHC) and quantification

Mice were deeply anesthetized with a ketamine/xylazine mixture and transcardially perfused with ice-cold PBS. Collected tissues were either immediately snap-frozen on dry ice for RNA or protein extraction, or post-fixed in freshly prepared 4% paraformaldehyde (PFA; Electron Microscopy Sciences) for histological analysis. Fixed tissues were post-fixed for an additional 2 hours at room temperature (RT), then washed in PBS (3 × 15 minutes), cryoprotected in 30% sucrose (Fisher Chemical) overnight at 4°C, and embedded in O.C.T. compound (VWR International). Serial coronal cryosections (12 μm) were cut using a Leica Cryostat (CM 1900-3-1) and stored at −80°C until use. For IHC, sections were air-dried at RT for 2 hours, then blocked in 10% donkey serum diluted in PBS containing 0.1% Triton X-100 for 1 hour at RT. Sections were incubated with primary antibodies overnight at 4°C, followed by PBS with 0.1% Tween-20 (PBST) washes and incubation with fluorescently labeled secondary antibodies for 2 hours at RT. DAPI was used as a nuclear counterstain. Images were acquired using a Nikon A1 confocal microscope. Confocal Z-stacks were obtained at 1 μm intervals (total thickness: 10 μm) and maximum intensity projections were generated for image analysis and quantification. To identify the cellular source of SERPINA3N, we quantified the percentage of SERPINA3N⁺ cells co-expressing specific lineage markers including GFAP (astrocytes), SOX10 (oligodendrocytes), and CD68 (activated microglia/macrophages). SERPINA3N^+^ cells that did not co-localize with any of these markers were categorized as “Other”. Quantification was performed on IHC images from at least three sections per animal (minimum n ≥ 4 animals per group, as indicated in the figure legends), and data are reported as the percentage of SERPINA3N⁺ cells in each category. Antibodies used for IHC are listed below: Goat polyclonal anti-Serpina3n (1:200, R&D System, Cat# AF4709; RRID:AB_2270116), Rabbit polyclonal anti-IBA1 (1:100, WAKO, Cat# 019–19741; RRID:AB_839504), Rabbit polyclonal anti-SOX10 (1:200, Abcam, Cat# ab27655;

RRID: AB_778021), Mouse monoclonal anti-GFAP (1:200, or 1:20000, Agilent, Cat# Z0334; RRID:AB_10013382), Rat monoclonal anti-CD68 (1:200, Bio-Rad, Cat# MCA1957; RRID: AB_3100585), Mouse monoclonal anti-APC (Ab-7) (CC1) (1:100, Millipore, Cat#OP80; RRID: AB_213434), Rabbit monoclonal anti-TCF4/TCF7L2 (C48H11) (1:100, Cell Signaling Technology, Cat#2569S; RRID:AB_2199816), Rabbit recombinant monoclonal anti-Stat3 (1:100, Cell Signaling, Cat# 4904, RRID:AB_331269), Rabbit recombinant monoclonal anti-pStat3 (Tyr705) (1:100, Cell Signaling, Cat# 9145; RRID:AB_2491009), Rabbit polyclonal anti-RFP (1:200, Rockland, Cat# 600-401-379; RRID:AB_2209751), Mouse monoclonal anti-Aβ1-16 (1:100, clone 6E10, BioLegend, Cat# 803001; RRID:AB_2715854), Rat monoclonal anti-I-A/I-E (1:100, BD Pharmingen, Cat# 556999; RRID:AB_396545), Mouse monoclonal anti-NeuN (1:100, Millipore, Cat# MAB377; RRID:AB_2298772), Rabbit recombinant monoclonal anti-phosphor-p65 (Ser536) (1:100, Cell Signaling Technology, Cat# 3033, RRID:AB_331284), Mouse monoclonal anti-CD45 (1:100, Thermo Fisher Scientific, Cat# 14-0451-82; RRID:AB_467251), Rabbit polyclonal anti-CD74 (1:100, Thermo Fisher Scientific, Cat# PA5-22113; RRID:AB_11157006), Chicken polyclonal anti-GFP (1:200, Abcam, Cat# ab13970; RRID:AB_300798), All 2^nd^ antibodies for IHC are from Jackson ImmunoResearch Laboratories.

#### RNA extraction, cDNA preparation, RT-qPCR, and primers

Total RNA was isolated using the RNeasy Lipid Tissue Mini Kit (QIAGEN) according to the manufacturer’s instructions, including on-column DNase digestion using the RNase-free DNase Set (QIAGEN) to eliminate genomic DNA contamination. RNA concentration and purity were assessed using a NanoDrop 2000 Spectrophotometer (Thermo Fisher Scientific). Complementary DNA (cDNA) was synthesized using the Omniscript RT Kit (QIAGEN). Quantitative real-time PCR (RT-qPCR) was carried out using the QuantiTect SYBR® Green PCR Kit (QIAGEN) on an Agilent MP3005P thermocycler. Gene expression was normalized to the internal control gene Hsp90, and relative expression levels were calculated using the 2^−ΔCt method: ΔCt = Ct(Hsp90) – Ct(target gene). Primer sequences used for RT-qPCR are listed below: Serpina3n (F/R, GCCTCGTCAGGCCAAAAAG/TGAACGTGTCAAGAGGGTCAA), Cxcl10 (CCCACGTGTTGAGATCATTG/CACTGGGTAAAGGGGAGTGA); C3 (AGCTTCAGGGTCCCAGCTAC/GCTGGAATCTTGATGGAGACGC); Il1b (GAAATGCCACCTTTTGACAGTG/CTGGATGCTCTCATCAGGACA); Ccl2 (CACTCACCTGCTGCTACTCA/GCTTGGTGACAAAAACTACAGC); Cd68 (TGTCTGATCTTGCTAGGACCG/GAGAGTAACGGCCTTTTTGTGA); Aldh1l1 (GCAGGTACTTCTGGGTTGCT/GGAAGGCACCCAAGGTCAAA); P2ry12 (CCCTGTGCGTCAGAGACTAC/CAAGCTGTTCGTGATGAGCC); Tnfa (TGTGCTCAGAGCTTTCAACAA/CTTGATGGTGGTGCATGAGA); iNos (CCCTTCAATGGTTGGTACATG/ACATTGATCTCCGTGACAGCC); Cd86 (GAGCGGGATAGTAACGCTGA/ GGCTCTCACTGCCTTCACTC); Cd206 (CTTCGGGCCTTTGGAATAAT/TAGAAGAGCCCTTGGGTTGA); Ym1 (CAGGTCTGGCAATTCTTCTGAA/GTCTTGCTCATGTGTGTAAGTGA); P21 (TCTTGCACTCTGGTGTCTGA/CTGCGCTTGGAGTGATAGAA); Hsp90 (AAACAAGGAGATTTTCCTCCGC/CCGTCAGGCTCTCATATCGAAT).

#### Protein preparation and Western blot

Tissues were lysed in N-PER Neuronal Protein Extraction Reagent (ThermoFisher), supplemented with protease and phosphatase inhibitor cocktail (ThermoFisher) and PMSF (Cell Signaling Technology). Lysates were incubated on ice for 10 minutes and centrifuged at 10,000 × g for 10 minutes at 4°C. Protein concentrations were determined using a BCA protein assay kit (ThermoFisher Scientific). Equal amounts of protein (30 μg per sample) were resolved by SDS-PAGE using AnykD Mini-PROTEAN TGX precast gels (Bio-Rad) and transferred to 0.2 μm nitrocellulose membranes (Bio-Rad) using the Trans-Blot Turbo Transfer System (Bio-Rad). Membranes were blocked with 5% BSA (Cell Signaling Technology) for 1 hour at room temperature and incubated overnight at 4°C with primary antibodies. After washing, membranes were incubated with appropriate HRP-conjugated secondary antibodies and visualized using Western Lightning Plus ECL (PerkinElmer). Band intensities were quantified using NIH ImageJ software. Antibodies used for Western blot are listed below: Goat polyclonal anti-Serpina3n (1:1000, R&D System, Cat# AF4709; RRID:AB_2270116), Rabbit polyclonal anti-IBA1 (1:1000, WAKO, Cat# 019–19741; RRID:AB_839504), Rabbit polyclonal anti-GFAP (1:1000, Millipore, Cat# MAB360; RRID:AB_11212597), Rat monoclonal anti-CD68 (1:1000, Bio-Rad, Cat# MCA1957; RRID: AB_3100585), Mouse monoclonal anti-β-actin (1:1000, Cell Signaling Technology, Cat# 3700; RRID:AB_2242334), Rabbit monoclonal anti-GAPDH (1:1000, Cell Signaling Technology Cat# 2118, RRID:AB_561053). All HRP-conjugated 2^nd^ antibodies are from Thermo Fisher Scientific.

#### RiboRag RNA immunoprecipitation

To analyze ribosome-bound transcripts in oligodendrocytes and astrocytes, Plp1-CreERT2 and Aldh1l1-CreERT2 transgenic lines were crossed with RiboTag mice (RRID:IMSR_JAX:029977). Adult mice received intraperitoneal tamoxifen injections once daily for five consecutive days. Starting 14 days after the final injection, mice were maintained on either CPZ diet or normal diet for 4 weeks prior to tissue collection. Brains were rapidly extracted and homogenized on ice using a Micro-Tube Homogenizer System in ice-cold homogenization buffer (10% w/v) containing: 50 mM Tris-HCl (pH 7.4), 100 mM KCl, 12 mM MgCl₂, 1% NP-40, 1 mM DTT, 0.1 mg/ml cycloheximide (Sigma), 200 units/ml RNasin (Promega), and complete protease inhibitor cocktail (MilliporeSigma), all prepared in RNase-free water. Homogenates were centrifuged at 10,000 × g for 10 minutes at 4°C to remove cellular debris. Supernatants were transferred to fresh microcentrifuge tubes kept on ice. A 10 µl aliquot was saved as the input control. For immunoprecipitation, 5 µl of anti-hemagglutinin (HA) antibody (Covance Anti-HA.11 Epitope Tag Antibody) or 1 µg of normal IgG control was added to the remaining 400 µl of cleared lysate. Samples were incubated for 4 hours at 4°C with gentle rotation. Protein A/G magnetic beads (Thermo Fisher Scientific, Cat. No. 88803; 100 µl per sample) were washed three times with homogenization buffer, equilibrated, added to the samples and incubated overnight at 4°C with rotation. The next day, samples were washed three times with high-salt buffer (50 mM Tris-HCl, 300 mM KCl, 12 mM MgCl₂, 1% NP-40, 1 mM DTT, 100 units/ml RNasin, 0.1 mg/ml cycloheximide, and 1:200 protease inhibitor in RNase-free water), with each wash lasting 5 minutes on a rotator in a cold room. After the final wash, magnetic beads were collected using a magnetic stand, and residual buffer was removed. Beads were resuspended in RLT plus lysis buffer, and RNA was extracted using the RNeasy Plus Micro Kit (Qiagen, Cat. No. 74034) following the manufacturer’s protocol.

#### Magnetic bead-assisted cell sorting (MACS)

Single-cell suspensions were prepared according to the instructions provided by Miltenyi Biotec. Mouse brains from both normal diet and cuprizone diet groups or CFA control, MOG control and MOG Serpina3n cKO groups were processed using the Adult Brain Dissociation Kit (Mouse and Rat) (Cat# 130-107-677, Miltenyi Biotec, Germany) in conjunction with the gentleMACS Dissociator (Cat# 130-093-235, Miltenyi Biotec, Germany). The mouse brains were cut into 0.5 cm pieces then transferred into a pre-heated gentleMACS C Tube (Cat# 130-093-237, Miltenyi Biotec, Germany). Enzymatic cell dissociation was initiated using 1,950 μL of enzyme mix 1 (Enzyme P and Buffer Z). The C Tube was attached upside down onto the sleeve of the gentleMACS Dissociator, and the brain tissue was dissociated using the appropriate gentleMACS program. After one rotation, 30 μL of enzyme mix 2 (Enzyme A and Buffer Y) was added to the C Tube, followed by two additional gentle rotations at 37°C. Upon completion of the program, the C Tube was detached, briefly centrifuged, and the sample collected at the bottom of the tube was filtered through a MACS SmartStrainer (70 μm) to remove cell clumps, thereby achieving a single-cell suspension. The MACS SmartStrainer was then washed with an additional 10 mL of D-PBS (Cat# 14287-080, Gibco) to ensure the collection of all cells. After centrifugation, the supernatant was gently removed, and the brain homogenate pellet was incubated separately with anti-CD11b MicroBeads (Cat# 130-093-634, Miltenyi Biotec, Germany) and anti-O4 MicroBeads (Cat# 130-094-543, Miltenyi Biotec, Germany). Followed by Fc receptor, anti-ACSA-2 MicroBeads (Cat# 130-097-678, Miltenyi Biotec, Germany) were used to separate astrocyte. The incubation was carried out for 15 minutes at 4°C in a refrigerator. The cells were then washed with 0.5% BSA/PBS buffer and centrifuged at 300 g for 10 minutes. After complete aspiration of the supernatant, the cell pellet was resuspended in 500 μL of 0.5% BSA/PBS buffer before proceeding to magnetic separation. To magnetic separate the glial cells magneteiclly, the LS MACS Column was positioned in the magnetic field of a MACS Separator (Cat# 130-090-976, Miltenyi Biotec) and rinsed with 3 mL of 0.5% BSA/PBS buffer. The cell suspension was applied to the LS Column, which was washed three times with 3 mL of 0.5% BSA/PBS buffer. Unlabeled cells were collected and combined with the flow-through, while magnetically labeled cells were immediately flushed out by firmly pushing the plunger into the column. Finally, 350 μL of Buffer RLT Plus containing β-mercaptoethanol (β-ME) was added to the cells, and the mixture was stored at −80°C.

#### RNA sequencing

For bulk RNA-seq, high-quality total RNA (RIN >7) was used to enrich mRNA using oligo(dT)-attached magnetic beads. The enriched poly(A) RNA underwent fragmentation, followed by first- and second-strand cDNA synthesis with dUTP incorporation to maintain strand specificity. The cDNA was end-repaired, 3′-adenylated, and ligated to bubble-shaped adapters before PCR amplification. The resulting PCR products were denatured, circularized using a bridged primer, and amplified via phi29 polymerase to form DNA nanoballs (DNBs). These DNBs were loaded onto patterned nanoarrays and sequenced on an MGI T7 system, producing paired-end 150 bp reads. After sequencing, raw reads were quality-checked and aligned to the Mus musculus reference genome (GCF_000001635.27_GRCm39) using HISAT or Bowtie2 (v2.3.4.3), then quantified by RSEM (v1.3.1). Differentially expressed genes (DEGs) were identified (e.g., DESeq2, threshold Q ≤0.05 or FDR ≤0.001). Subsequent analyses, including principal component analysis, correlation, and GO/KEGG enrichment, were conducted to interpret gene expression patterns.

For scRNA-seq, single cell suspension was prepared from approximately 50–100 mg of flash-frozen tissue via homogenization, filtration, and low-speed centrifugation. After confirming >80% cell viability (trypan blue), each suspension was processed on the 10x Genomics Chromium platform, generating GEM (Gel Bead in Emulsion) droplets with unique barcodes and UMIs. Reverse transcription was carried out within each droplet, followed by cDNA purification, amplification, fragmentation, end-repair, adapter ligation, and PCR indexing. Libraries were optionally circularized and amplified into DNA nanoballs before sequencing on the DNBSEQ G400 system (PE28+100, ∼350M reads per sample). Resulting reads were aligned to the Mus musculus reference genome (refdata-gex-mm10-2020-A) using Cell Ranger (v5.0.1) ^81^. Seurat (v3.2.0)^82^ was employed for downstream quality filtering (<200 genes per cell, top 15% mitochondrial fraction, and doublet exclusion via DoubletDetection^83^. A set of 2,000 highly variable genes was used for principal component analysis (PC=15), followed by dimensionality reduction with UMAP. Cluster marker genes were identified using the FindAllMarkers function (logfc.threshold>0.25, min.pct>0.1, Padj≤0.05), and SCSA^84^ was utilized to assist in cell-type annotation.

#### Gene set score calculation

For each defined gene set, expression values were first transformed via log₂(expression+1) to stabilize variance and mitigate the impact of highly expressed genes. We used the previously published strategy to calculate gene set scores ^18^. In cases where all genes were anticipated to move in a consistent direction, the sample score was calculated as the arithmetic mean of the log-transformed values. Where both up- and downregulated genes were involved, each gene was assigned a +1 or –1 weight based on its presumed direction of regulation, and the final score was obtained by averaging these weighted, log-transformed values. If baseline correction was needed, the mean gene set score of control samples was subtracted from each test sample. All analyses were conducted in R (v4.4.0).

#### Primary microglia culture and SERPINA3N treatment

Primary cortical cell cultures were prepared from postnatal day 0-2 mouse pups. Mouse pups were quickly decapitated with sterile scissors and then the heads immediately were transferred into ice-cold saline solution. The meninges were carefully removed under an anatomical microscope, and the cortices were then mechanically dissociated with surgical instruments and transferred with a 10 mL pipette to a sterile 50 mL conical tube. After centrifugation, cortices tissues were added with papain dissociation kit (Cat# LK003176, Worthington, USA) supplemented with DNase I (250 U/mL, Cat# D5025, Sigma Aldrich, USA) and D-(+)-glucose (0.36%, Cat# 0188, AMRESCO, USA), and incubated in 36 °C/10% CO_2_ chamber for 90 min. Digested tissues were pipetted up and down with a sterile Pasteur pipette and then flushed to the cell strainer. Strained cells were centrifuged at x 300 g for 10 min. Cells were then plated on poly-D-lysine (PDL, Cat# A003-E, Millipore, USA)-coated T-75 flask (Cat# 430641U, Corning, USA) by adding DMEM/F12 (Cat# 1196092, Thermo Fisher, USA) with 10% heat-inactivated fetal bovine serum (FBS, Cat# 12306-C, Sigma Aldrich, USA) and 1% penicillin/streptomycin (P/S, Cat# 15140122, Thermo Fisher, USA). The plates were incubated in a 5% CO_2_ incubator at 37 °C for 10-14 days to grow a confluent mixed glial population. After mixed glial culture was completely confluent, a mechanic shaking method was used to isolate microglia (180 rpm at 37 °C without CO_2_ for 2 hr). After shaking, culture media containing microglia in the T-75 flask was transferred to 50 mL tubes. Microglia were washed with PBS and cell count was proceeded using a hemocytometer with Trypan Blue exclusion. Next, microglia were resuspended with DMEM/F12 containing 10% FBS, 1% P/S, and microglia supplement (Cat# 1952, ScienCell, USA) and seeded appropriately onto 6-well cell culture plates. All culture plates were incubated in a 37°C with 5% CO_2_ incubator and the media was changed every other day after seeding. Primary microglia at 14 DIV were stimulated with LPS (Cat# L2880, *E. coli* O55:B5, Sigma Aldrich, USA) (10ng/ml) for 24 hours with or without recombinant SERPINA3N (50 ng/mL, Cat# 4709-PI, R&D System, USA).

#### Data collection and statistics

Data collection and quantification were performed by lab members blinded to mouse genotype and treatment. Both male and female mice were included in all experiments. Data are presented as mean ± standard error of the mean (s.e.m.) unless otherwise indicated individually. Scatter dot plots were used throughout the manuscript, with each dot representing a single mouse or one independent experiment. Data normality was assessed using the Shapiro-Wilk test. For comparison between two groups, an unpaired two-tailed Student’s t test was used. The degrees of freedom are reported as t(df) in figure legends. For comparisons involving more than two groups, a one-way ANOVA followed by Tukey’s multiple comparisons test was performed. The F-statistic and associated degrees of freedom are reported as F_(DFn, DFd)_, where DFn and DFd represent the numerator and denominator degrees of freedom, respectively. Brown-Forsythe test was used to evaluate variance equality across multiple groups. All statistical analyses and data visualizations were conducted using GraphPad Prism version 8.0. A *p*-value of less than 0.05 was considered statistically significant. Significance is indicated in figures as follows: ^∗^p < 0.05, ^∗∗^p < 0.01, ^∗∗∗^p < 0.001; “ns” denotes not significant (*p* > 0.05).

## DATA AVAILABILITY

All RNA-seq data used in the study were submitted to Gene Expression Omnibus (GEO) database and are publicly available. GSE identifications were assigned as follows: GSE287324 (Suppl Table 1 and Suppl Table 4), GSE287323 (Suppl Table 2); GSE287320 (Suppl Table 3); GSE287310 (Suppl Table 6, Suppl Table 7, Suppl Table 8, and Suppl Table 9); GSE287313 (Suppl Table 10); GSE287309 (Suppl Table 12). Other data are available upon request.

## Acknowledgements

We thank the funding agencies of NIH (R21NS125464, R01NS123080, R01NS123165, R01NS134887 to FG, R01NS069726 to SD) and Shriners Hospitals for Children (85101-NCA-22, 85113-NCA-23 to FG, 85410-NCA-24 to YW, 84312-NCA-24 to MZ, 84331-NCA-24 to JP) for supporting the work.

